# Multiomics Reveals *IGF-1* c.258A > G Reduces Triacylglycerol Containing Medium-Chain Fatty Acid in Milk

**DOI:** 10.64898/2026.09.16.752017

**Authors:** Shuo Zheng, Jiayuan Fang, Yi Li, Xingyu Xiao, Tong Su, Qinchuan Lv, Yunyun Cheng, Yuanyuan Xu, Linlin Hao

**Author notes:** Corresponding authors: (Y Xu), (L Hao). Equal contribution (Shuo Zheng, Jiayuan Fang).

## Abstract

Fatty acid composition of milk triacylglycerol (TAG) is important for neonatal growth. We previously identified a synonymous mutation widely distributed in human populations, *IGF-1* c.258A > G, which decreased milk TAG content in gene-edited mice. So far, the acyl chain composition of affected TAGs, underlying molecular mechanism, and cross-species conservation remain unclear. Here, integrated transcriptomic and metabolomic analyses of mouse mammary glands revealed that this mutation downregulates IGF-1 expression and elevates lipoprotein lipase (LPL) activity, and potential associations between IGF 1 and peroxisome proliferator-activated receptor γ (PPARG), fatty acid synthase (FASN) and lipin 1 (LPIN1). Association analysis in 295 Bama pigs carrying this gene polymorphism showed reduced milk lipid content in sows; further, correspondingly lower litter weight was significantly associated with the genotype. Integrated metabolomic profiling of mouse mammary glands and porcine milk demonstrated insufficient production of lauric and myristic acids. Accordingly, supplementing these two fatty acids restored TAG content. Mechanistically, IGF-1 promotes triacylglycerol-containing medium-chain fatty acid (MCFA-TAG) synthesis by activating the PPARG-FASN/LPIN1 axis, whereas increased LPL activity indirectly suppresses this axis through long-chain fatty acid accumulation. Together, our results reveal the molecular mechanism whereby this mutation affects milk lipid synthesis, providing insights into the regulation of MCFA-TAG synthesis.

## Introduction

Milk lipids, a unique energy source for neonates, delivers >50% of milk’s caloric intake. TAG, the major component of milk lipids, account for up to 98% of total milk lipid content; fatty acid composition and TAG distribution are critical for infant neurodevelopment, bone health, and intestinal homeostasis [1]. In particular, MCFA-TAGs not only improve fat absorption and support brain development in neonates [2,3], but also modulate gut microbial activity and enhance hepatic fatty acid oxidation in neonate animals [4,5]. There are considerable differences in milk lipid composition among different mammalian species and their genotypes. Specifically, ∼80% of African American mothers carry *FADS* genetic variants associated with elevated arachidonic acid levels, compared with only 45% of European American mothers [6]. Similarly, previous studies found that *MC4R* p.298G > A had an association with colostrum lipid composition in Landrace and Large White pigs [7]. Despite these observations, research into genetic variants linked to milk lipids is still limited, especially concerning the specific genetic locus associated with milk MCFA-TAG-related traits and their regulatory mechanisms.

IGF-1 can induce endoplasmic reticulum biogenesis in mammary epithelial cells [8], a process tightly linked to *de novo* lipid synthesis. Additionally, genetic polymorphisms at other loci within *IGF-1* are associated with milk lipid content in Holstein cows [9]. In our previous studies, we identified a synonymous mutation c.258A > G in the highly conserved *IGF-1* coding region. This mutation exhibits variable allele frequencies across human ethnic groups and reduces IGF-1 mRNA stability [10,11], and decreased milk TAG content through a mechanism associated with lipoprotein lipase (LPL) in gene-edited mice. While the molecular mechanisms by which this mutation regulates TAG metabolism are clear, its impact on population genetics remains uncertain.

*De novo* TAG synthesis in the mammary gland primarily utilizes endogenously synthesized fatty acids as substrates, subsequently esterified under catalysis of multiple lipogenic enzymes [12]. Peroxisome proliferator-activated receptor γ (PPARG), a key transcriptional regulator in this process, drives *de novo* fatty acid synthesis and conversion of phosphatidic acid to diacylglycerol by activating the expression of target genes, including fatty acid synthase (FASN) and lipin 1 (LPIN1) [13,14]. LPL hydrolyzes TAG to generate LCFAs [15], including the potent PPARG agonists eicosapentaenoic acid and conjugated linoleic acid, whereas increased LCFAs suppress PPARG transcriptional activity [16,17]. Compared with LCFAs, MCFAs, including lauric acid and myristic acid, weakly activate PPARG within physiological ranges [18–20]. The acid-labile subunit of the insulin-like growth factor, which binds to IGF-1 and modulates its bioavailability, is known to activate PPARG signaling [21]; nonetheless, the functional association between IGF-1, LPL, and PPARG during milk lipid synthesis remains unclear.

To clarify how *IGF-1* c.258A > G affects milk lipid synthesis and its cross-species conservation, we employed three experimental models: *IGF-1* c.258A > G gene-edited mice with a clean genetic background, 295 Bama pigs with human-like lactational physiology carrying this polymorphism [22,23], and variant-harboring human mammary epithelial MCF-10A cells. Integrated multiomics of mouse mammary glands revealed associations between IGF-1 and PPARG, FASN, and LPIN1 in the glycerolipid metabolism. Association analysis in 295 sows linked the mutant genotype to reduced milk lipid content and lower litter weight. Cross-species lipidomic profiling confirmed insufficient lauric and myristic acids. In mutant MCF-10A cells, supplementation with these fatty acids restored TAG content. Mechanistically, IGF-1 promotes MCFA-TAG synthesis via the PPARG-FASN/LPIN1 axis, whereas mutation-induced LPL elevation indirectly suppresses this axis through LCFA accumulation. This study highlights the functional significance of synonymous mutations in lactation genetics, providing insights into the regulation of mammary MCFA-TAG synthesis.

## Results

### Decreased IGF-1 expression and milk lipid content in the mammary gland of *IGF-1* c.258 A > G mice

The highly conserved *IGF-1* harbors a synonymous mutation at the c.258 locus, which presents genetic polymorphism in human populations and diverse pig breeds (Figure S1). Our previous research showed altered mammary lipid metabolism in *IGF-1* c.258A > G mice [24]. Here, the mutation reduced IGF-1 mRNA and protein levels in the mammary gland (**Figure 1**A and B), consistent with our previous finding that this synonymous mutation decreases mRNA stability. Cross-species relative synonymous codon usage analysis confirmed that GCG is a low-frequency codon compared to GCA, without differential m A modification (Figure S2). Next, to further assess its effect on milk lipid content, we first measured milk yield and found a 20% reduction in Ho mice at lactation day 2 (L2d) (Figure 1C). Concurrently, TAG concentration was significantly decreased in both mammary glands and milk of Ho mice (Figure 1D and E). Milk lipids are synthesized by mammary epithelial cells (MECs), assembled into lipid droplets (LDs), and secreted as milk fat globules (MFGs) [25]. Although LDs were successfully synthesized and secreted in MECs of both genotypes, the number of LDs in the mammary lumen was markedly decreased in Ho mice (Figure 1F). Moreover, despite comparable MFG sizes, the number of MFGs in Ho mice decreased by 56% (Figure 1G and H). More importantly, independent of pup genotype, pups fostered by Ho dams exhibited a lower average body weight than those nursed by WT dams, without pup mortality in both groups (Figure 1I), indicating that the reduced pup body weight was due to maternal milk lipid content rather than pup genotype or survival issues. Together, these findings demonstrate that *IGF-1* c.258A > G leads to decreased IGF-1 expression and lower TAG content in the mammary gland.

**Figure 1.**
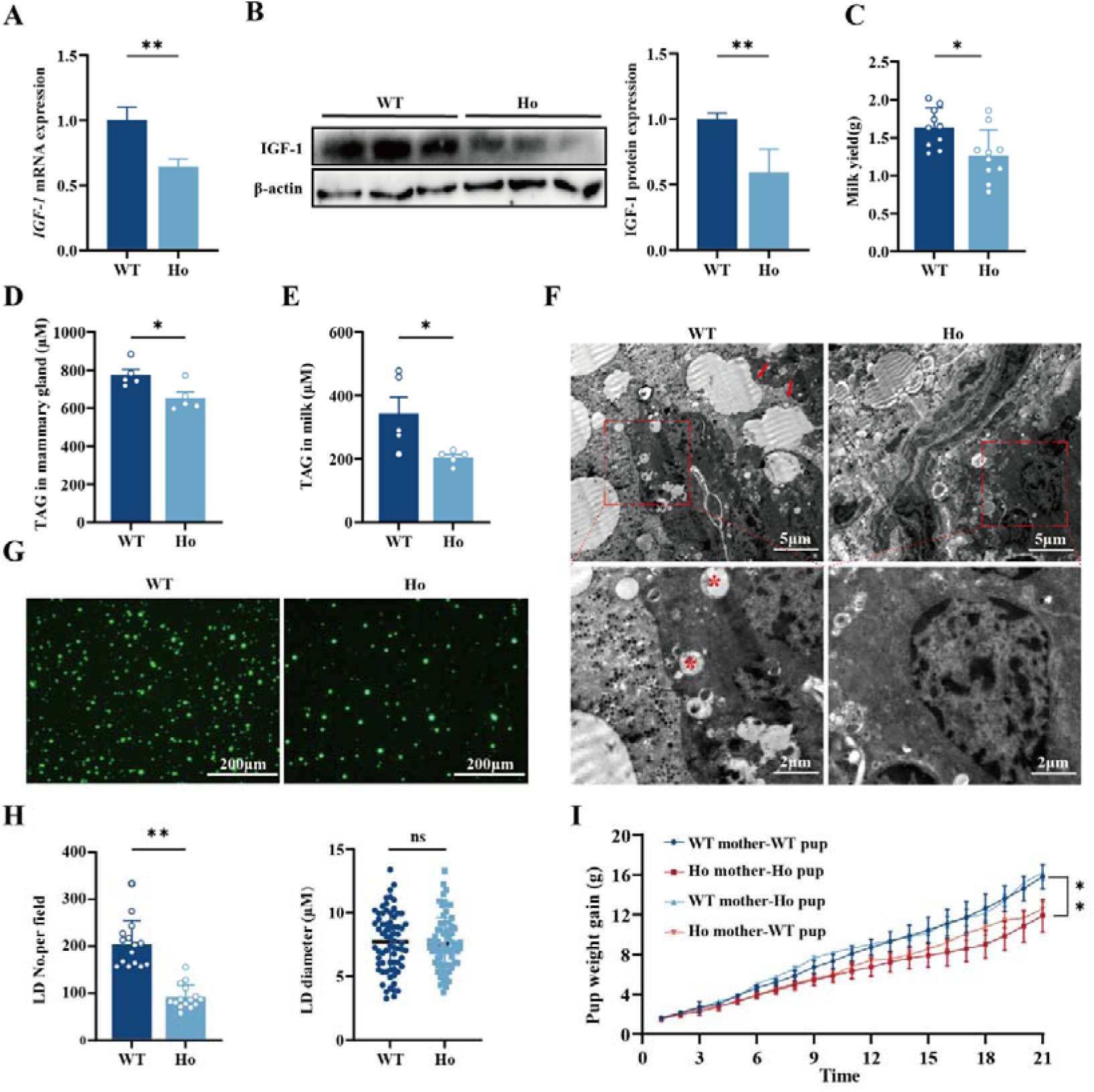
***IGF-1* c.258 A > G mice exhibit lower milk lipid content and pup weight A. and B.** IGF-1 mRNA and protein expression of mice at L2d, lactation day 2. **C.** Milk yield in L2d mice (n = 10 for each genotype). **D. and E.** TAG, Triacylglycerol concentration in mammary glands and milk of mice at L2d. **F.** Electron microscopic images of mammary gland sections in L2d mice. Asterisks (_*_) indicate cytoplasmic lipid droplets and arrowheads MFGs, milk fat globules. **G. and H.** Bodipy staining of MFGs in L2d mice and quantification. **I.** Body weight of cross-nursed pups during lactation. Data are presented as mean ± SEM (n = 5 dams for each cross-fostering group, 8 pups per dams). Data are presented as mean ± SEM. \**P* < 0.05, \*\**P* < 0.01, ns *P* > 0.05.

### Inhibited *de novo* TAG synthesis in the mammary gland

Milk lipid synthesis depends on normal mammary gland development and systemic lipid metabolism; accordingly, mammary ductal morphogenesis and the expression of lipid metabolism-related genes were evaluated. No significant differences in the number of mammary ductal branches and terminal end buds were observed (Figure S3A–D). Moreover, compared with Ho mice, circulating IGF-1 and its major transport protein IGFBP3 [26] levels were elevated in WT mice, without changes in the expression of lipolysis-related genes in adipose tissue and the liver (Figure S3E and F). Only PRL mRNA expression in the pituitary was decreased in Ho mice (Figure S3G). Thus, the decreased milk lipid content was not significantly associated with mammary gland development or mammary exogenous lipid metabolism.

To dissect how reduced IGF-1 impaired mammary milk lipid synthesis, we performed transcriptomic sequencing on the mammary glands of WT and Ho mice at L2d. Principal component analysis (PCA) showed a distinct separation between groups, indicating that the mutation altered the transcriptomic profile of the mouse mammary gland. A total of 392 differentially expressed genes (DEGs), including 153 upregulated and 239 downregulated genes, were identified (Figure S4A; Table S1). Gene ontology (GO) enrichment analysis revealed enriched DEGs in the lipid metabolism and lipase activity, which directly correlate with impaired milk lipid synthesis (**Figure 2**A). Kyoto Encyclopedia of Genes and Genomes (KEGG) pathway analysis demonstrated that downregulated DEGs were involved in sphingolipid and fatty acid synthesis (*GLA*, *CERS6*, *FASN*, *ACACA*), compromising TAG and MFG membrane synthesis. Meanwhile, DEGs in glycerolipid and glycerophospholipid metabolism (*PLPP4*, *GPAT*, *LPIN1*) disrupted TAG and MFG assembly. In addition, the dysregulated arachidonic acid metabolism (*Cyp2c29*) affected lipid-derived signaling (Figure 2B–C). Additionally, LPL expression in the mammary gland was upregulated (Figure S5A), accompanied by a concurrent ∼1.5-fold increase in LPL enzymatic activity (Figure S5B). The mammary gland consistently showed elevated free fatty acid concentrations as shown in Figure S5C.

**Figure 2.**
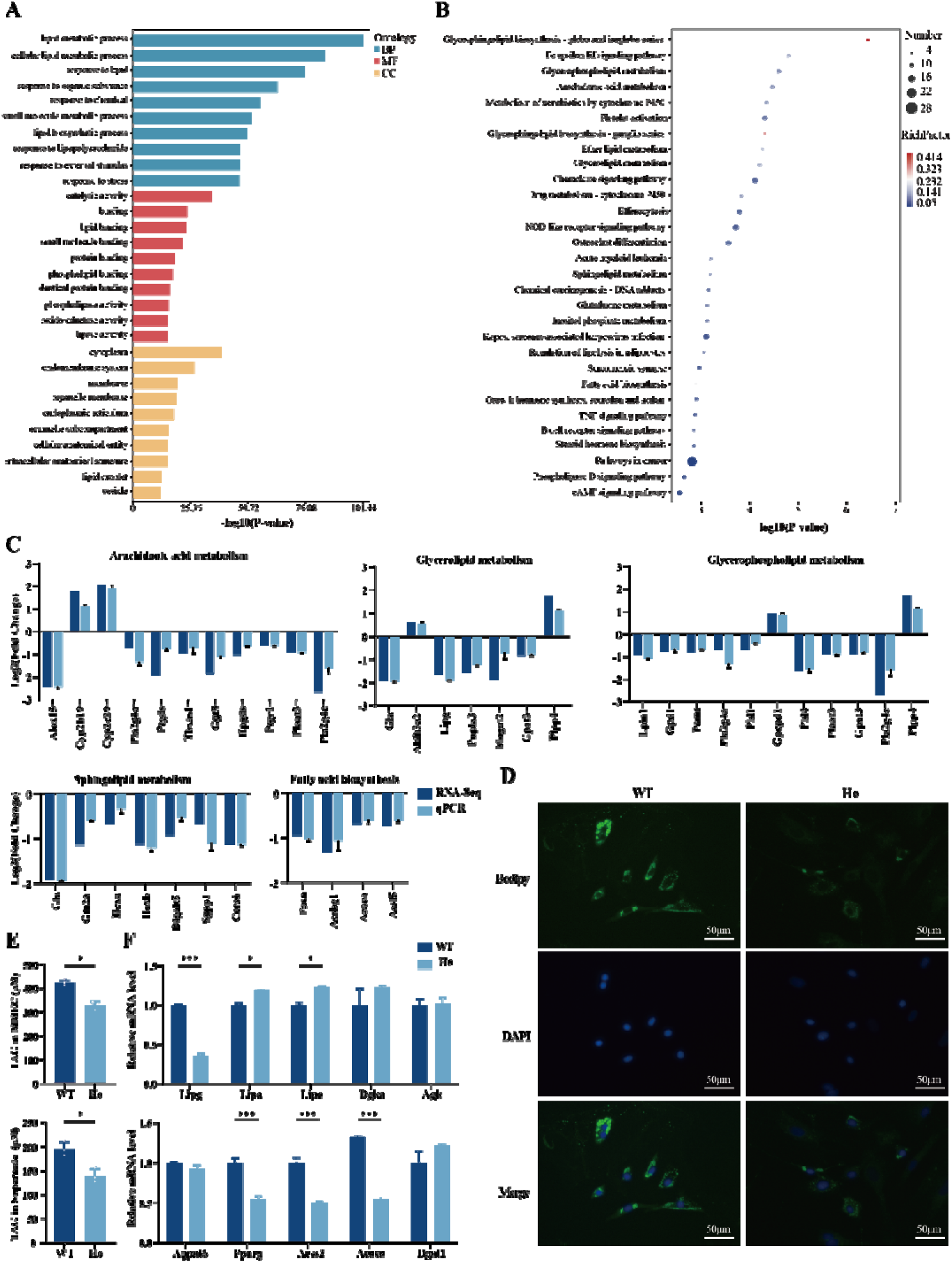
***IGF-1* c.258A > G only alters milk lipid metabolism in the mammary gland A.** Top 10 enriched GO, gene ontology pathways of DEGs, differentially expressed genes of each subgroup according to *P*-value. **B.** Top 30 enriched KEGG, Kyoto Encyclopedia of Genes and Genomes pathways of DEGs according to *P*-value. **C.** Validation of DEGs involved in different lipid metabolism pathways by qPCR. **D.** Bodipy staining of LDs, lipid droplets in primary mouse MECs, mammary epithelial cells. **E.** TAG concentration in MECs and cell culture supernatants. **F.** mRNA expression of genes involved in TAG synthesis and degradation in MECs. Data are presented as mean ± SEM. * *P* < 0.05, *** *P* < 0.001.

Further validation confirmed that the number of LDs was reduced in primary MECs from Ho mice (Figure 2D and S6); in addition, decreased TAG concentration in cell and cell culture supernatants (Figure 2E) was consistent with findings in mice. Furthermore, *ACACA*, a key gene involved in fatty acid synthesis, and *ACSS2*, a critical gene mediating fatty acid activation [27], were both downregulated. In contrast, mRNA levels of *LIPA* and *LIPE*, which are involved in triglyceride hydrolysis [28,29], were upregulated (Figure 2F). This reduction in milk lipid content was achieved by specifically inhibiting *de novo* milk lipid synthesis within the mammary gland.

### PPARG/FASN/LPIN1 are key targets in IGF-1 regulated milk lipid synthesis

To elucidate the mechanism whereby reduced IGF-1 suppresses milk lipid synthesis, transcriptomic and lipidomic analyses were added to mouse mammary gland data. DEGs and differential lipid metabolites (DLMs) were co-enriched in 15 pathways, including lipid metabolism-related pathways, such as glycerophospholipid and glycerolipid metabolism (**Figure 3**A). Since fatty acid synthesis primarily provides precursors for these two pathways [30], we further integrated these data with pathway-specific DEGs and DLMs to construct a regulatory network for milk lipid synthesis (Figure 3B; Table S2). Notably, these three pathways were co-enriched in five triglyceride metabolites, four of which exhibited a downregulation trend, consistent with the reduced milk lipid content. Subsequently, protein-protein interaction (PPI) network analysis of the DEGs identified in the MEC regulatory network revealed potential interactions between PPARG, FASN and LPIN1, and IGF-1 (Figure 3C). Among them, PPARG is a key transcriptional regulator that directly modulates the expression of milk lipid synthesis-related genes *FASN* and *LPIN1* [31], both consistently downregulated in Ho mice (Figure 3D). Consequently, PPARG, FASN and LPIN1 exhibited downregulated expression in association with IGF-1. These are crucial genes involved in suppressing milk lipid synthesis.

**Figure 3.**
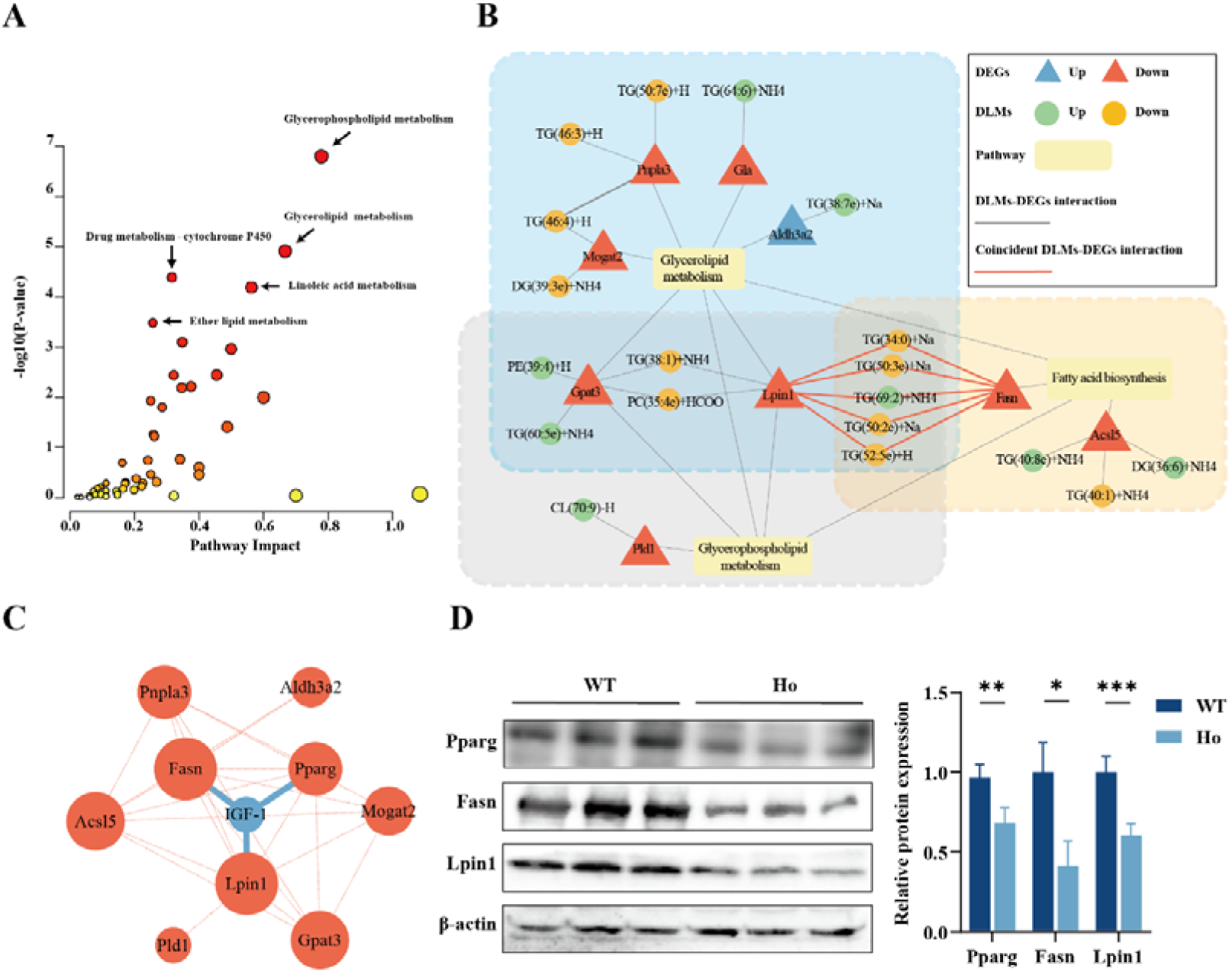
Reduced IGF-1 impairs milk lipid synthesis by inhibiting PPARG, FASN, and LPIN1 expression. **A.** KEGG pathways co-enriched by DEGs and DLMs, differential lipid metabolites according to *P*-value. **B.** Regulatory network of milk lipid synthesis and metabolism, including lipid metabolites and genes significantly regulated by *IGF-1* c.258A > G. **C.** PPI, Protein-protein interaction network between IGF-1 and proteins related to milk lipid metabolism. The larger the circular node, the greater the number of its interacting partners. Lines connecting different circular nodes represent the interactions; blue lines represent interactions with IGF-1. **D.** Expression of proteins interacting with IGF-1 in L2d mice. Data are shown as mean ± SEM. \**P* < 0.05, \*\**P* < 0.01, \*\*\**P* < 0.001.

### *IGF-1 c.258A > G* reduces milk lipid content in sows and correlates with weaning litter weight in piglets

While inhibited milk lipid synthesis has been demonstrated in *IGF-1* c.258A > G mice, rodents exhibit distinct mammary gland structure and milk lipid composition from other mammals. Thus, Bama pigs with genetic polymorphisms were selected to further investigate the effects of this mutation on milk lipid content and composition. Three genotypes at the *IGF-1* c.258 locus, AA (wild-type, WT), AG (heterozygous, He), and GG (homozygous, Ho), were identified in 295 Bama sows (**Figure 4**A). This locus exhibited valid polymorphism (PIC = 0.3539, HW*P*-val > 0.05; **Table 1**).

**Figure 4.**
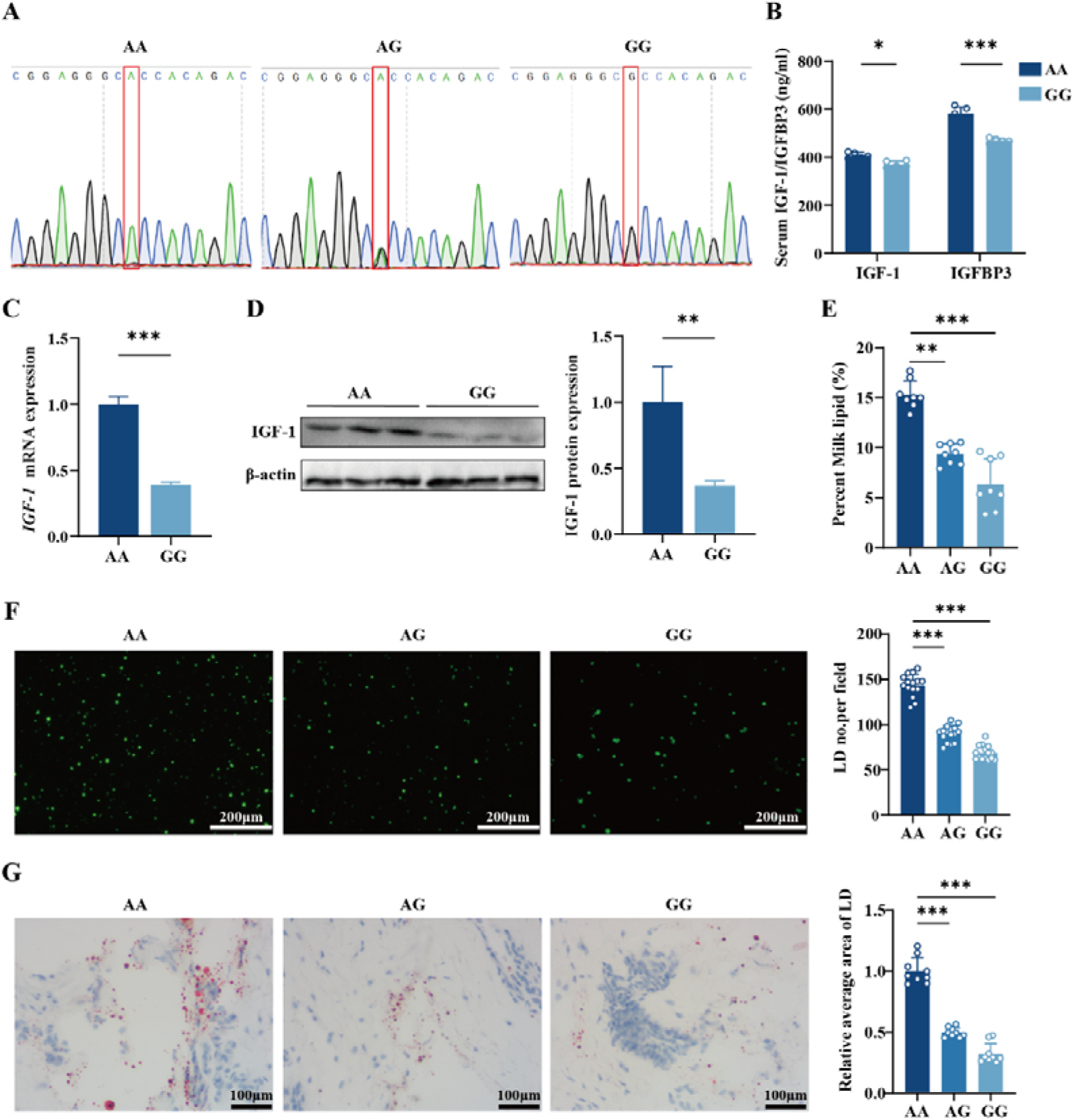
***IGF-1* c.258 A > G decreases IGF-1 expression and milk lipid content in Bama sows A.** Sequencing diagram of the three genotypes at the *IGF-1* c.258 locus. **B.** Circulating IGF-1 and IGFBP3 levels in L2d Bama sows. **C. and D.** IGF-1 mRNA and protein expression analysis of Bama sows at L2d. **E.** Milk lipid content in L2d sows (n = 8 for each genotype). **F.** Bodipy staining and quantification of MFGs in the milk of L2d sows. **G.** Oil Red O staining and quantification of LD area in the mammary glands of L2d sows. Data are shown as mean ± SEM. \**P* < 0.05, \*\**P* < 0.01, \*\*\**P* < 0.001.

**Table 1.** Genetic diversity parameters of the IGF-1 c.258 locus in Bama sows

| locus | Location | Position | Genotype | Genotype frequency | Allele | Allele frequency | Ho | He | HWP-val | PIC |
| --- | --- | --- | --- | --- | --- | --- | --- | --- | --- | --- |
| c.258<br>A > G | Exon<br>4 | 5:8183<br>0812 | AA | 0.3966 | A | 0.6424 | 0.5085 | 0.4915 | 0.4485 | 0.3539 |
|  |  |  | AG | 0.4915 |  |  |  |  |  |  |
|  |  |  |  |  | G | 0.3576 |  |  |  |  |
|  |  |  | GG | 0.1119 |  |  |  |  |  |  |

Note: He, gene heterozygosity; Ho, gene homozygosity; HWP-val, *P*-value of Hardy–Weinberg Equilibrium; PIC, Polymorphism information Content.

Consistent with the finding that *IGF-1* c.258A > G can reduce mammary IGF-1 expression and milk lipid content in mice, AG and GG sows exhibited decreased IGF-1 expression and serum IGF-1 and IGFBP3 levels (Figure 4B–D), and ∼1.5-fold lower milk lipid content compared to AA sows (Figure 4E). Meanwhile, a greater number of LDs and MFGs were observed in the mammary gland and milk of AA sows than in AG and GG sows (Figure. 4F and G). Furthermore, an association analysis revealed that sows and piglets with the AA genotype had higher milk yield and litter weight weaned than those with AG and GG genotypes (**Table 2**). Importantly, the preweaning mortality rate tended to be higher in GG piglets than in AA piglets **(Table 3)**. In general, *IGF-1* c.258A > G decreases milk lipid content by suppressing IGF-1 expression in both Bama pigs and mice, thereby affecting neonatal weight gain.

**Table 2.** Association between the *IGF-1* c.258 locus and reproductive and lactation performance of sows and piglets

| Genotype | Milk yield(kg) | Litter size | Live births | Still births | Litter weight(kg) | Litter weight weaned(kg) |
| --- | --- | --- | --- | --- | --- | --- |
| AA (117) | 136.660±4.668<sup>a</sup> | 10.309±0.278 | 9.125±0.253 | 1.184±0.176 | 10.700±0.641 | 78.254±1.743<sup>a</sup> |
| AG (145) | 123.383±3.521<sup>b</sup> | 9.963±0.270 | 8.867±0.245 | 1.097±0.171 | 11.094±0.404 | 72.059±1.691<sup>b</sup> |
| GG (33) | 109.616±5.053<sup>c</sup> | 10.557±0.426 | 9.183±0.387 | 1.374±0.270 | 11.066±0.418 | 64.703±2.668<sup>c</sup> |
| *P*-value | 1.343E-5 | 0.202 | 0.451 | 0.523 | 0.132 | 1.271E-4 |

*Note:* Milk yield (kg) = piglet ADG × litter size × lactating days × 4 [79]; a,b within the same column, means with different superscript letters indicate differences at $P < 0.05$ ; data are presented as mean ± SEM.

**Table 3.** Effects of *IGF-1* c.258, A > G on mortality rates (%) of sow-reared piglets during lactation

| Genotype | D0 | D7 | D14 | D21 | D28 |
| --- | --- | --- | --- | --- | --- |
| AA | 0.00 | 3.679±8.894 | 3.914±8.982 | 4.056±9.052 | 4.056±9.052 |
| AG | 0.00 | 3.759±10.491 | 3.954±10.562 | 4.154±10.749 | 4.212±10.749 |
| GG | 0.00 | 4.432±7.709 | 5.844±8.953 | 6.119±9.192 | 6.178±10.049 |
| *P*-value | — | 0.467 | 0.194 | 0.247 | 0.273 |

*Note:* Data are presented as mean ± SEM.

### Decreased MCFA-TAG content in the milk of *IGF-1* c.258A > G sows

Given that *IGF-1* c.258A > G induces dysregulation of mammary lipid metabolism, lipidomic analysis was performed to further evaluated its effects on milk lipid composition and distribution in populations harboring this polymorphism. PCA revealed significant differences in the expression profiles of lipid metabolites between WT and Ho groups (Figure S2B). A total of 697 lipid metabolites were identified in milk. These included glycerolipids, sphingolipids, glycerophospholipids and other classes, with TAGs being the most abundant (**Figure 5**A; Table S4). Among all lipid classes, the levels of several lipid subclasses were significantly altered due to *IGF-1* c.258A > G. Specifically, relative TAG levels were decreased. In contrast, as products of lipid catabolism, fatty acid and lysophospholipid levels including lyso-phosphatidylcholine, lyso-phosphatidylethanolamine, and lyso-phosphatidylinositol, were relatively increased [32]. Similarly, the levels of phosphatidylcholine, phosphatidylethanolamine, and phosphatidylinositol — major components of the MFG membrane [33], as well as ceramide phosphate, ganglioside, hexCer, sphingomyelin, and sulfatide, core components of lipid rafts [34], were decreased (Figure 5B). Further visualization of 247 DLMs revealed a general trend toward downregulation of TAG, which was not only the lipid subclass with the highest number of DLMs but also the metabolites experiencing the most substantial fold changes (Figure 5C). This indicates that TAG downregulation is the key factor underlying the reduction in milk lipid content triggered.

**Figure 5.**
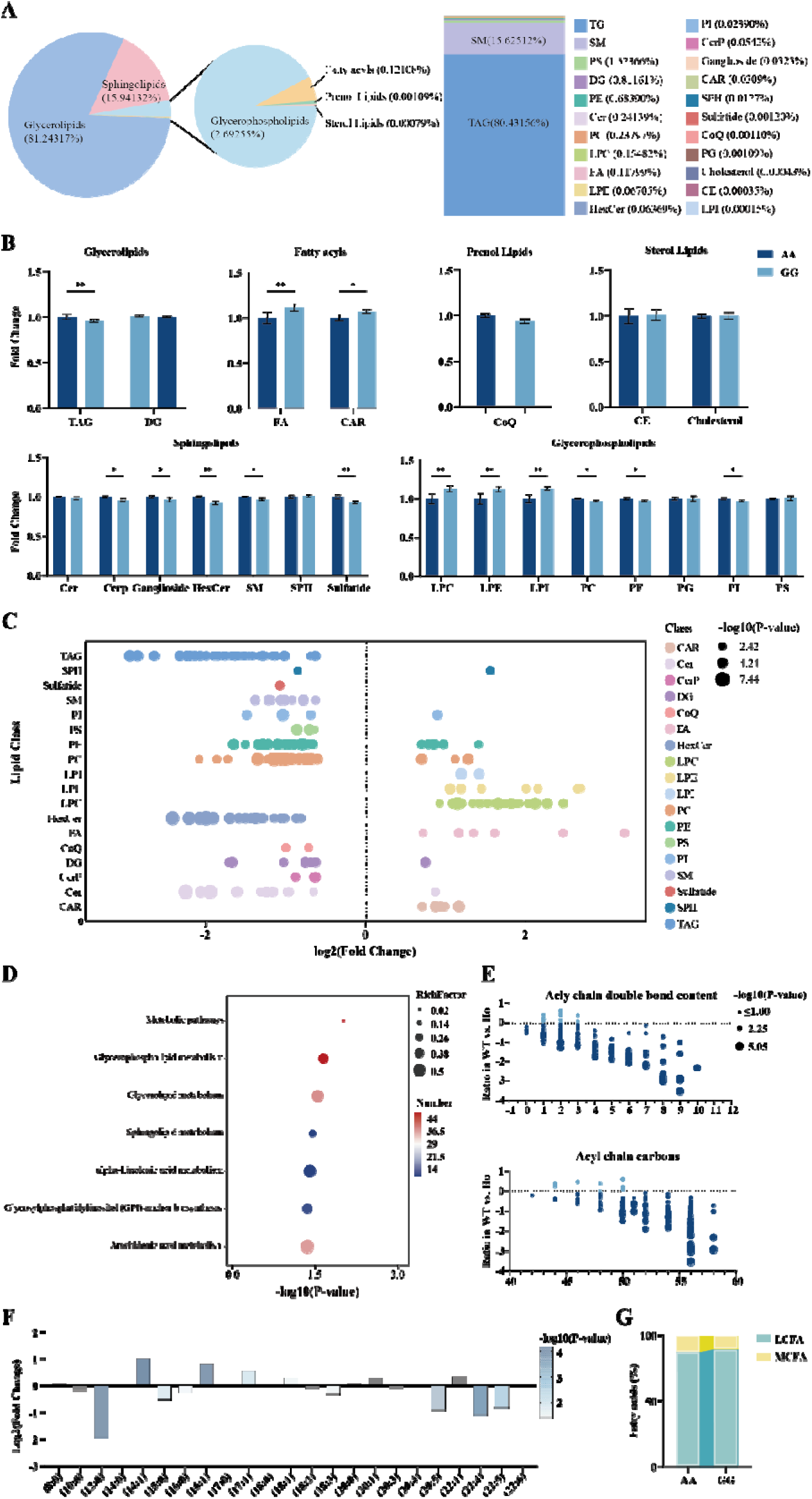
***IGF-1* c.258A > G alters milk lipid composition A.** Distribution of milk lipid classes (left) and subclasses (right) in L2d Bama sows. **B.** Intensity fold change of the overall lipid composition. **C.** Distribution of intensity fold changes across lipid classes. Each dot represents a lipid species within a specific class; dot size indicates statistical significance. **D.** KEGG pathways enriched by DLMs according to *P*-value. **E.** Comparison of TAG profiles between WT and Ho types. Each dot represents a distinct TAG, arranged along the x-axis based on the total number of double bonds (top) or number of carbon atoms (bottom) in the acyl chain. Dot size is proportional to the *P*-value. Only lipid species with *P < 0.05* are displayed. **F.** Intensity distribution of individual fatty acyl chains associated with TAGs. Bar transparency is proportional to the significance value, expressed as -log10 (*P*-value). Gray bars represent those with *P* > 0.05. **G.** Comparative analysis of medium-chain and long-chain acyl chain contents in TAGs. Data are presented as mean ± SEM. \**P* < 0.05, \*\**P* < 0.01.

In addition, KEGG enrichment analysis confirmed that glyceride, glycerophospholipid, and sphingolipid metabolism are the most significantly differentially regulated lipid metabolic pathways (Figure 5D). Notably, *IGF-1* c.258A>G reduced the content of TAGs with high double-bond content (>3 insaturations) and high acyl-chain carbon numbers (>50 carbons) (Figure 5E). We also observed a significant decrease in the relative levels of MCFAs, whereas LCFAs remained unchanged (Figure 5F and G). In addition, LCFAs exhibited increased relative abundance (Figure S7). In glycerophospholipids, lysophospholipids showed elevated relative levels of fatty acyl chains, while most phosphatidyl lipids, particularly long-chain fatty acyl chains (>18 carbons) indicative of MFG membrane synthesis, displayed reduced relative abundance (Figure S8A–F). Similarly, reduced levels of sphingolipid subtypes with long acyl chains (≥18 carbons) compromise lipid droplet stability (Figure S9A–D).

### Decreased levels of lauroyl-CoA and myristoyl-CoA suppress TAG synthesis

Combined multiomics analyses of sow milk and mouse mammary gland consistently demonstrated that reduced IGF-1 impairs TAG synthesis (Figure 3A and 5D). TAG synthesis, a regulated metabolic process involving multiple rate-limiting enzymes, begins with FASN-catalyzed fatty acid production, followed by LPIN1-mediated integration and assembly to ultimately form TAG. PPARG plays a key role as a transcriptional regulator in this process, as shown in **Figure 6**A. *De novo* TAG synthesis mainly relies on MCFAs and partial LCFAs in the mammary gland. Notably, most LCFAs are derived from exogenous substrates transported to the mammary gland, whereas no significant alterations are observed in systemic lipid metabolism [35]. More importantly, the mutation significantly reduced the relative levels of MCFAs (Figure 5G) and phosphatidic acid levels in the mammary gland (Figure S10), whereas total DAG levels showed no significant change (Figure 5B). Thus, we focused on the dynamic changes in MCFAs. Combined lipidomic profiling of porcine milk and mouse mammary gland demonstrated that the mutation decreased the abundance of TAGs conjugated with lauroyl-CoA and myristoyl-CoA (Figure 6B). Therefore, we treated MCF-10A cells harboring the *IGF-1* c.258A > G variant (Ho type, Figure S11A) with 100 μM lauric acid and 200 μM myristic acid at safe and effective concentrations (Figure S11B). MCFA supplementation promoted LD accumulation, increased TAG concentration, and upregulated LPIN1, PPARG, and FASN expression (Figure 6C-E). These findings indicate that reduced IGF-1 expression suppresses FASN - and LPIN1 - mediated TAG synthesis, thereby reducing the MCFA levels essential for TAG synthesis.

**Figure 6.**
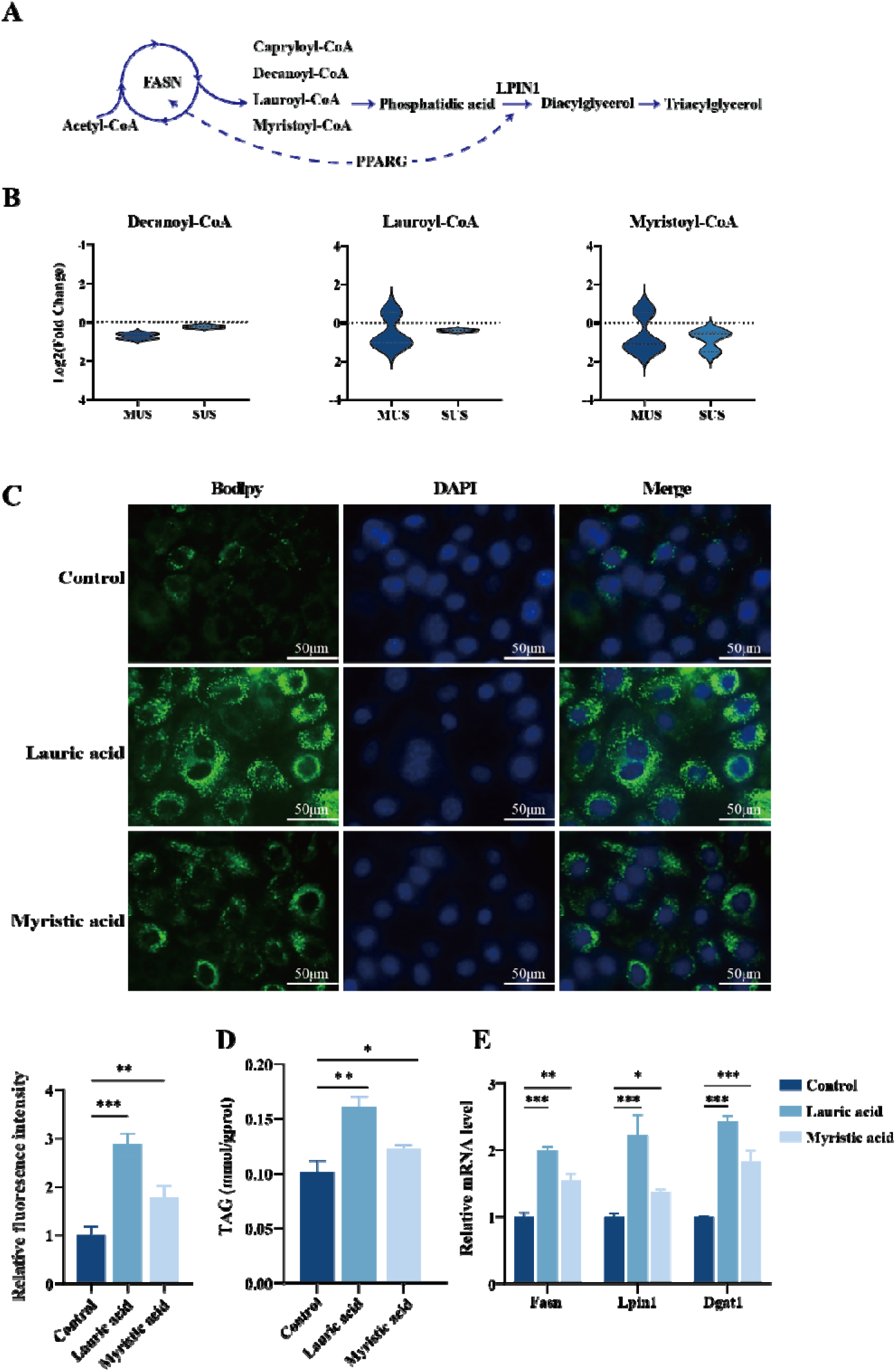
Chain fatty acyl content for TAG synthesis decreases in both sow milk and mouse mammary gland. **A.** Schematic diagram of *de novo* TAG synthesis. **B.** Abundance of fatty acyl-CoA species among TAGs. **C**. Bodipy staining of LDs (up) and relative fluorescence intensity (down) in MCF-10A cells treated with lauric acid and myristic acid. **D**. TAG concentration in MCF-10A cells treated with lauric acid and myristic acid. **E**. The mRNA expression of PPARG, FASN, and LPIN1 in MCF-10A cells treated with lauric acid and myristic acid. Data are presented as mean ± SEM. \**P* < 0.05, \*\**P* < 0.01.

### Milk lipid synthesis is suppressed via the IGF-1-PPARG-FASN/LPIN1 axis

Integrated transcriptome and lipidome analyses of the mouse mammary gland revealed that reduced IGF-1 expression suppressed PPARG and then downregulated milk lipid synthesis-related genes, including *FASN* and *LPIN1*. Further, treatment with rosiglitazone (Rosi) increased TAG concentration and LD accumulation (Figure S12A and B), and upregulated FASN and LPIN1 protein levels (Figure S12C and D). To further investigate whether IGF-1 regulates milk lipid synthesis via the PPARG-FASN/LPIN1 axis, IGF-1 signaling was either activated or blocked using an IGF-1 overexpression vector (OE-IGF1) or the IGF-1R inhibitor Picropodophyllin (PPP), respectively (Figure S13A and B). IGF-1 overexpression increased TAG content and LD accumulation (**Figure 7**A and B), upregulated PPARG, FASN, and LPIN1 protein levels, and partially antagonized the inhibitory effect of GW9662 on this axis (Figure 7C). Conversely, PPP treatment significantly reduced TAG content and LD formation (Figure 7D and E), and downregulated their protein expression of these genes (Figure 7F). Rosi treatment partially alleviated the inhibitory effect induced by PPP (Figure 7F). These results indicate that IGF-1 promotes milk lipid synthesis through activation of the PPARG-FASN/LPIN1 axis. Furthermore, since LPL hydrolyzes lipoproteins at the surface of MEC to release LCFA, Ho cells were supplemented with an LCFA mixture containing elevated species and proportions in Ho mammary glands to mimic the downstream effects of increased LPL activity (Figure S13C). This treatment reduced TAG content and LD formation while suppressing the PPARG-FASN/LPIN1 axis. Combined PPP and LCFA treatment further enhanced this inhibitory effect (Figure S13D–F). These findings suggest that IGF-1 indirectly regulates milk lipid synthesis through the LPL-LCFA pathway.

**Figure 7.**
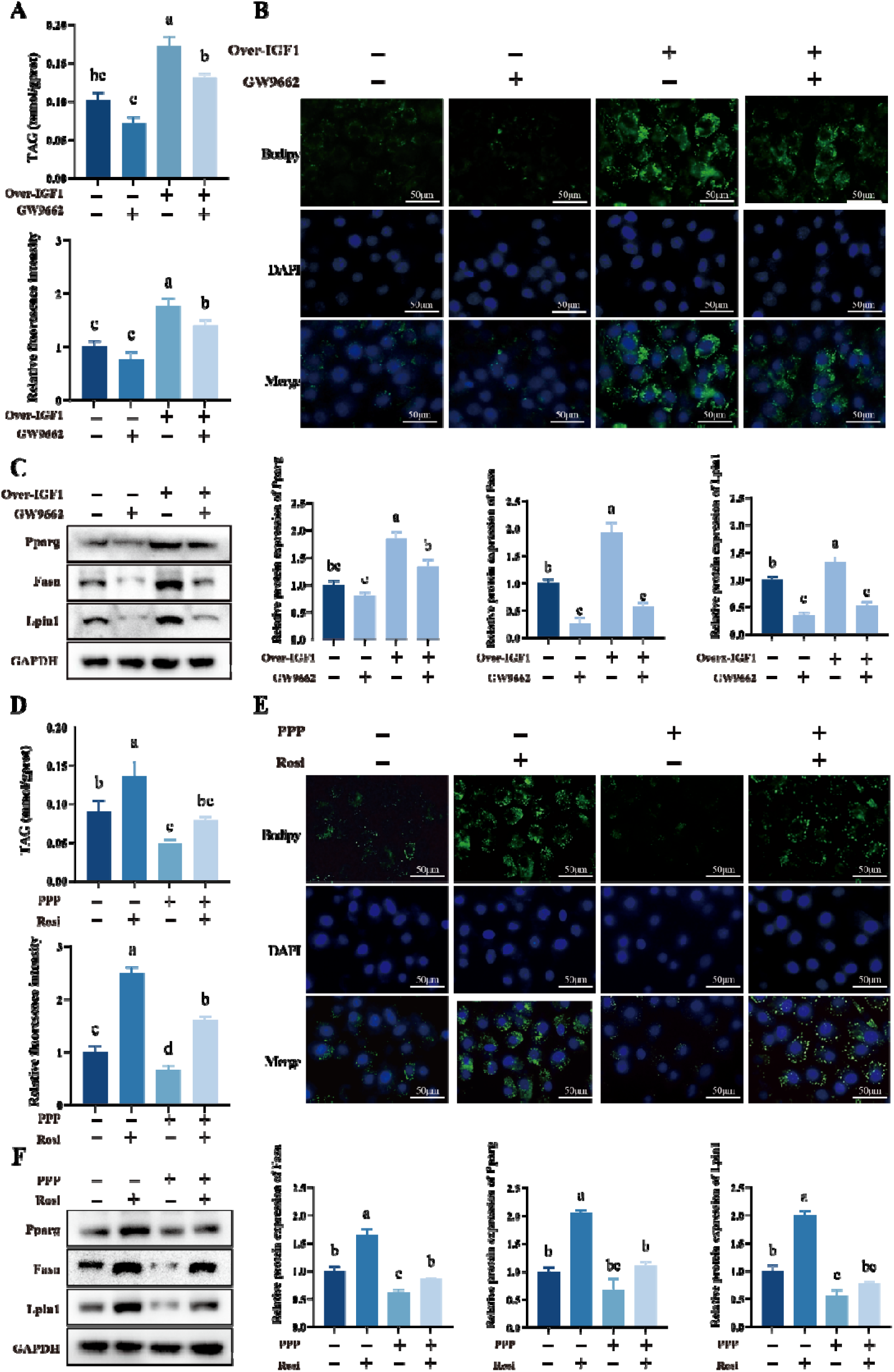
IGF-1-PPARG-FASN/LPIN axis regulates MCFA-TAG synthesis in the mammary gland A–C. TAG content (A), bodipy staining of LDs (right) and relative fluorescence intensity (left) (B) and protein expression levels of PPARG, FASN, and LPIN1 (C), in MCF-10A cells overexpressing IGF-1 and treated with GW9662. **D–F.** TAG content (D), bodipy staining of LDs (right) and relative fluorescence intensity (left) (E), and protein expression of PPARG, FASN, and LPIN1 (F) in MCF-10A cells treated with PPP, Picropodophyllin and Rosi, rosiglitazone. Data are presented as mean ± SEM. Different letters indicate significant differences among groups (*P* < 0.05).

## Discussion

Milk lipids provide the primary energy and nutrient for neonates, and their content and composition are determined by genetic factors [36,37]. Synonymous mutations, long regarded as "functionally silent," have not been sufficiently explored in milk lipid metabolism. In this study, we first demonstrated that *IGF-1* c.258A > G synonymous mutation downregulates IGF-1 expression, inhibits MCFA-TAG synthesis via PPARG-FASN/LPIN1, and ultimately reduces milk lipid content. These findings highlight the non-negligible role of synonymous mutations in milk lipid metabolism.

Synonymous mutations modulate phenotypic traits through mechanisms such as epitranscriptomic modifications or altered translational efficiency [38,39]. Cheng and Wang confirmed that *IGF-1* c.258A > G affects IGF-1 expression through altered mRNA and protein stability [11,40]. Here, we demonstrated that the mutation downregulates IGF-1 expression in the mammary gland of gene-edited mice and Bama pigs through altered codon usage bias. Consistent with previous research, elevated circulating IGF-1 can promote milk secretion, and upregulation of local IGF-1 in the mammary gland can enhance endoplasmic reticulum lipid synthesis [41,42]. Of note, most IGF-1 functional mutations are concentrated in regions associated with skeletal development [43]. Only a few association analyses have indicated a correlation between *IGF-1* mutations and milk fatty acids [44]. Nevertheless, the underlying regulatory mechanisms remain unclear. The coding region of the *IGF-1* gene exhibits over 80% sequence homology across different mammalian species [45]. This conservation pattern is analogous to that of the *FUS* gene, which exerts conserved regulatory functions across multiple species [46], implying that the functional *IGF-1* c.258A > G may also be involved in lactation across multiple mammalian species.

Efficient synthesis of milk lipids relies on the coordinated effects of normal mammary gland development, sufficient supply of fatty acid precursors, and appropriate expression of lipid synthesis-related genes [47]. Studies in liver-specific IGF-1 knockout mice confirm that a 75% reduction in circulating endocrine IGF-1 does not impair mammary ductal development [48], but directly impairs lactation [42]. In this study, mutant mice showed only a 4% decrease in circulating IGF-1, with normal mammary development and systemic lipid metabolism. Combined with the finding that IGF-1 overexpression in MECs directly promotes TAG synthesis, these results demonstrate that the decreased local mammary IGF-1 expression induced by the mutation is the core cause of impaired milk lipid synthesis [49–51]. Meanwhile, given the mammary gland’s cellular diversity, milk lipid synthesis is uniquely controlled by specialized secretory cell clusters [52]. Notably, the expression of lipid synthesis-related genes, including *ACACA*, *ACSS2*, and *PPARG* [31], was downregulated in Ho MECs. These results indicate that *IGF-1* c.258A > G specifically regulates *de novo* lipid synthesis in the mammary gland.

*De novo* milk lipid synthesis in the mammary gland utilizes fatty acids as the core substrate: on the one hand, it synthesizes TAG as the main energy component; on the other hand, it participates in glycerophospholipid assembly to form the MFG membrane [53]. Integrated transcriptomic and lipidomic analyses of the mammary gland revealed that *LPIN1*, involved in glycerolipid and glycerophospholipid metabolism, and *FASN*, involved in fatty acid metabolism, were both correlated with five TAGs. Among them, four were downregulated, consistent with the decreased TAG content observed in the mammary glands of Ho mice. Further analysis confirmed that IGF-1 has potential interactions with PPARG, FASN, and LPIN1. Previous studies have shown that IGF-1 can interact with PPARG to regulate embryonic development [54], and with LPIN1 and FASN in breast cancer cells [55,56]. Notably, we previously demonstrated that the mutation led to increased LPL expression. Excessive LPL activity hydrolyzes TAGs to release LCFAs, consistent with the increased free fatty acid content detected in mutant mammary glands. When LCFA production exceeds the esterification capacity of MECs, lipotoxic intermediates accumulate, exerting negative feedback inhibition on the expression of the aforementioned genes [57–59]. However, an interaction between IGF-1 and these genes has not been reported in MECs.

This study in the Bama pig model further confirmed that the mutation reduces the milk lipid content of sows and weaned litter weight of piglets, with an increasing tend in preweaning mortality of piglets nursed by GG sows. Concurrently, milk lipidomic analysis delineated specific changes in milk lipid composition. Consistent with the findings in the mouse model, significant alterations were observed in the profiles of glycerolipids and glycerophospholipids in milk. Reduced IGF-1 disrupted the balance between lysophospholipids and glycerophospholipids, which may lead to abnormal MFGM structure and reduced efficiency of lipid absorption in piglet intestines [60]. Reduced IGF-1 also decreased the relative levels of sphingolipids, another major component of polar lipids, which have varied biological functions, including nervous system development, intestinal maturation, and preventing infection [61–63]. Interestingly, the alteration patterns of glycerolipids and sphingolipids in porcine milk mirrored those seen in mouse mammary glands. However, glycerophospholipids in the mammary gland remained unchanged. This could be because glycerophospholipids in tissues primarily maintain the membrane integrity of acinar cells and cell signaling, necessitating greater stability of their overall content [64]. In contrast, glycerophospholipids in milk vary with lactation stage [65,66]. Additionally, the higher relative abundance of LCFAs in milk supports the observed increase in mammary LPL activity. More importantly, the TAG content in the milk of GG sows decreased, accounting for >80% of the total detected milk lipids. The mutation significantly decreased MCFA-TAG content in milk. Specifically, lauroyl-CoA and myristoyl-CoA showed a consistent decline across both animal models. MCFAs are crucial for providing quick energy to a piglet’s intestine, and their reduced levels severely hinder the efficiency of milk lipid utilization [67]. Dietary glyceryl laurate supplementation in lactating sows increases piglet weaning weight [68]. Consistent with this finding, exogenous MCFA treatment rescued TAG levels in mutated MECs, confirming that MCFA deficiency impaired milk lipogenesis.

As weak agonists, MCFAs freely cross the plasma membrane in unesterified form to access the PPARG ligand-binding domain, activating lipogenic genes, including FASN and LPIN1. In contrast, LCFAs with higher binding affinity are preferentially esterified and sequestered in LDs, restricting nuclear activation. Mutation-induced LCFA overaccumulation exceeds mammary epithelial esterification capacity, causing lipotoxic intermediate buildup that ultimately suppresses PPARG transcriptional activity [58,69]. This is consistent with the observed increase in mammary FFA and decrease in milk lipid content in mutant mice. Similarly, combined treatment with PPP and excess LCFA in mutant MECs further suppressed the PPARG-FASN/LPIN1 axis. Furthermore, IGF-1 overexpression activated the PPARG-FASN/LPIN1 signaling axis, promoted TAG accumulation, and partially reversed the GW9662-induced inhibition of lipogenesis. Conversely, blockade of IGF-1 signaling by PPP suppressed this axis and reduced TAG content.

Collectively, these findings indicate that IGF-1 promotes milk lipid synthesis by activating the PPARG-FASN/LPIN1 axis, whereas the observed LCFA accumulation and decreased milk lipid content in mutants suggest that loss of IGF-1 signaling may indirectly exacerbate lipotoxicity-mediated PPARG suppression by modulating LPL activity. Given the variable allele frequencies of *IGF-1* c.258A > G across human populations and the high evolutionary conservation of *IGF-1*, MCF-10A cells only confirm the cross-species relevance of this axis. Future studies should validate the association between this genotype and human breast milk lipid profiles, as well as its impact on infant growth, and employ isotope-labeled internal standards for absolute quantification of key lipids.

## Conclusion

This study showed that *IGF-1* c.258A > G synonymous mutation downregulates IGF-1 expression, thereby inhibiting MCFA-TAG synthesis through the PPARG-FASN/LPIN1 signaling axis and ultimately decreasing milk lipid content. Our results deepen the understanding of IGF-1 function in lactation, fill a critical gap in synonymous mutation-mediated regulation of milk lipid traits, and provide a theoretical basis for understanding the mechanism of MCFA-TAG synthesis in mammary glands.

## Materials and methods

### Animals and phenotype collection

Wild type ICR mice (WT) were purchased from Changsheng Bio-technology Co. Ltd (Liaoning, China). *IGF-1* c.258 A>G ICR mice (Ho) were previously constructed in the laboratory [40] and bred for >5 generations to ensure stable inheritance. WT and Ho mice were used to ensure uniform genetic backgrounds and functional readouts in this gene-edited model. All mice were housed under SPF conditions (22±1°C, 50±5% humidity, 12 h light/dark cycle) with free access to sterilized feed and water. Milk yield was measured at L2d using the weigh-suckle-weigh method [70]. Briefly, pups were separated from dams for 4 h, weighed collectively, returned to the dams for 1 h of suckling, and reweighed. The difference in weight before and after suckling was considered as milk yield per hour. The effect of milk on pup growth was demonstrated through the method for cross-fostering [71]. To eliminate litter size effects, litters were standardized to eight pups on the first day. A total of 20 litters were used for cross-fostering, with five litters per cross-fostering group: (i) WT pups nursed by WT dams (WT mother-WT pup), (ii) WT pups nursed by Ho dams (WT mother-Ho pup), (iii) Ho pups nursed by WT dams (Ho mother-WT pup), and (iv) Ho pups nursed by Ho dams (Ho mother-Ho pup). The corresponding body weights and survival status of the pups were recorded for 21 days.

A total of 295 healthy Bama sows with parities ranging from 1 to 7 were selected based on the previously estimated heritability of milk traits from Bama Original Breeding Fragrant Pig Agriculture and Animal Husbandry Industrial Co, Ltd (China) to ensure a statistical power >80%. To evaluate natural allelic variation and additive genetic effects in an association analysis framework, three genotypes were retained in the pig population study. These sows were raised under standardized conditions, including food and water *ad libitum*, a 12 h light/dark photoperiod, and ambient temperature maintained at 20±2°C. Genotyping at the *IGF-1* c.258 locus classified sows into three: WT (AA), He (AG), and Ho (GG). Litter size, live births, stillbirths, and birth litter weight were recorded after farrowing. Weights and mortality status of piglets were recorded at farrowing, day 0, 7, 14, and 21 of lactation and weaning. The above phenotypes were collected from 295 sows for association analysis.

### Animal sample collection

Mouse dams (WT and Ho, n = 5) were separated from their pups for 1 h, followed by an injection of 10 units of oxytocin (O3251, Sigma, St. Louis MO) to induce milk letdown at L2d. Milk samples were collected from the 4^th^ pair of mammary glands [72], and blood, livers, and pituitary samples were collected at L2d. Mammary gland samples were collected on pregnancy day 18 (P18d) and L2d to assess mammary gland development.

Part ear samples of 295 sows were collected for genotyping. Milk samples for milk composition analysis were collected at L2d from the 3^rd^ to 6^th^ pair of teats of healthy second parity Bama sows (WT/He/Ho, n = 8), given the absence of milk cisterns, short teat length, and limited milk volume per gland in Bama pigs. Milk samples for lipidomic analysis were collected under identical screening criteria (WT, n = 7 and Ho, n = 6). Mammary gland samples of six sows (WT and Ho, n = 3) with similar parity were collected after euthanasia at L2d.

### Genotyping and association analysis

Genomic DNA was obtained from ear tissue samples using the Genomic DNA Kit (DP304, Tiangen, China). The primer (F: GGCCCCTCTGCATTTGATTTG R: GCACAGTACATCTCCAGCCT) used to amplify 700 bp containing c.258 regions of the IGF-1 gene was designed by reference to the porcine reference genome sequence (NC_010447.5). PCR products were sequencrf using an ABI3730XL DNA Analyzer (Applied Biosystems, Carlsbad MA) and genotyped with CHROMAS v2.23 software. Genetic diversity parameters, including genotype frequency, allele frequency, gene heterozygosity (He), gene homozygosity (Ho), Hardy–Weinberg equilibrium *P*-value (HW*P-*val), and polymorphism information content (PIC), were calculated using PopGene (v1.32). Correlations involving the *IGF-1* c.258 locus were analyzed using analysis of covariance (ANCOVA) implemented in SPSS (v26.0). The linear mixed model was structured as follows:

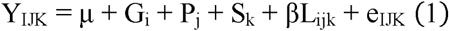

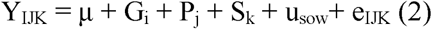

Where Y_IJK_ is the observed phenotypic value, μ the overall population mean, G_i_ the fixed effect of genotype (i = AA, AG, GG), P_j_ the fixed effect of parity (j = parity 1, parity 2–4, parity 5–7); S_k_ the fixed effect of farrowing season (k = spring, summer, autumn, winter), L_ijk_ the number of still births as a continuous covariate, and β the regression coefficient for this covariate, u_sow_ the random effect of sow, e_IJK_ the random error. Model (1) was used for milk yield, litter weight, litter weight weaned, litter size, live births, and still births. The covariate L was not included in models for litter size, live births, and still births. Model (2) was used for preweaning piglet mortality, where Y represents the mortality rate calculated as (live births − live piglets at the indicated time point) / live births × 100%. Pairwise comparisons among genotypes were performed using the least significant difference test. All phenotypic data are presented as least squares mean ± standard error of the mean (SEM).

### Primary mouse MEC isolation, culture, and identification

The 4^th^ pair of mammary glands from L2d mice were washed in D-Hank’s solution containing penicillin and streptomycin. They were then fragmented and digested with a mixed enzyme solution at 37°C for 1 h. Subsequently, the tissue suspension was subjected to three rounds of differential centrifugation at 1,500 ×g to discard fibroblasts and adipocytes and treated with 0.64% ammonium chloride (diluted in PBS) to lyse erythrocytes. Finally, cells were resuspended in basal DMEM/F12 medium with 500 ng/mL hydrocortisone (H0888, Sigma), 10 ng/mL epidermal growth factor (PHG0311, Invitrogen, Irvine CA), 100 U/mL penicillin/streptomycin (P1400, Solarbio, China), and 10% FBS (Gibco, New Zealand). MECs were sequentially fixed, permeabilized, and blocked before incubation with a primary antibody against cytokeratin 18 (1:200, 10830-1-AP, Proteintech, China) and a secondary antibody conjugated to Alexa Fluor 488 (1:4000, SA00013-2, Proteintech). Finally, for cell purity verification, nuclear staining was performed using DAPI (G1012, Servicebio, China), and images acquired with a Nikon Ti2 fluorescence microscope (Tokyo, Japan).

### MCF-10A cell culture and treatment

Human mammary epithelial cells (MCF-10A) were cultured in a complete medium consisting of DMEM/F12, 20 ng/mL epidermal growth factor, 0.5 μg/mL hydrocortisone, 10 μg/mL insulin, and 1% non-essential amino acids. To induce lactogenic differentiation, cells were treated with 5 μg/mL human prolactin (HY-NP198, MCE, NJ) [73]. For experiments, cells were transfected with 3 μg of pcDNA3.1-IGF1 or pcDNA3.1-empty vector using Lipofectamine™ 2000 (Thermo Fisher, Waltham MA) at 80% confluence. At 24 h post-transfection, cells were treated with 10 μM rosiglitazone (HY-17386, MCE), a PPARG agonist, and lysed for total RNA and protein extraction. In some experiments, cells were also treated with MCFA and LCFA. To mimic the elevated LCFA microenvironment in Ho mouse mammary glands, an LCFA mixture was prepared based on lipidomic analysis showing significantly increased total LCFA levels in Ho mice. It contained oleic acid (HY-N1446, MCE), arachidonic acid (HY-109590, MCE), and α-linolenic acid (HY-N0728, MCE) at a 4:1:0.3 ratio, and was added to a medium containing fatty acid-free BSA at gradient concentrations for 24 h. As determined by the triacylglycerol concentration assay, the mixture at final concentrations of 400 μM oleic acid, 100 μM arachidonic acid, and 32 μM α linolenic acid was verified to reduce the triacylglycerol content, which was selected for subsequent experiments. To verify whether MCFA supplementation could rescue milk lipid synthesis in mutant cells, lauric acid (HY-Y0366, MCE) and myristic acid (HY-N2041, MCE) were used at 100 and 200 μM at the highest non-cytotoxic concentrations, respectively, as determined by the CCK-8 cytotoxicity assay (IV08, Invigentench, Carlsbad CA). Briefly, CCK-8 solution was added at a 10:1 ratio (DMEM/F12 : CCK-8), incubated for 1 h, and the absorbance read at 450 nm. Complete medium served as negative control and CCK-8-containing medium without cells as blank.

Cell viability (%) = [(OD(sample) - OD(blank)) / (OD(negative) - OD(blank))] × 100%.

### Milk lipid content analysis

The milk lipid percentage from sows (WT/He/Ho, n = 6) was measured using the creamatocrit method [74]. Briefly, 75 µL milk sample was loaded into a hematocrit tube and centrifuged. Lipid layer height and total sample height were measured using a vernier caliper, and milk lipid percentage calculated as (lipid layer height / total sample height) × 100.

TAG concentration in mouse milk (WT and Ho, n = 5) or mammary epithelial cells (MEC and MCF-10A) was measured using a Triglyceride Content Assay Kit (E1003, Applygen, China).

### BODIPY staining

Mouse and sow’s milk were diluted in PBS and stained with Bodipy493/503 (D2191, Thermo Fisher). MECs were first fixed with 4% paraformaldehyde for 20 min, before staining with Bodipy 493/503 and DAPI (G1012, Servicebio) at room temperature, and photographed using an inverted fluorescence microscope (Nikon). MFG number and size were quantified using ImageJ software in ≥5 random fields per sample.

### Histology, H&E, and Oil red O staining

Mammary gland samples from mice were treated with an ethanol solution and embedded in paraffin, whereas those from sows were treated with a sucrose solution and embedded in OCT. Both samples were sectioned into 6 µm slices. Mouse mammary gland sections were stained with hematoxylin-eosin (G1120, Solarbio), whereas sow mammary gland sections were stained with Oil Red O (G1261, Solarbio). All images were captured under an inverted microscope (Nikon).

### Transmission electron microscopy

Mammary gland samples from L2d mice were sequentially fixed in glutaraldehyde and osmic acid, followed by staining in an aqueous uranium acetate solution.

Subsequently, samples were dehydrated and embedded. After ultrathin sectioning, LDs and MFGs in the mammary glands were imaged using a Tecnai Spirit Biotwin transmission electron microscope at 120 kV.

### Whole-mount staining and morphologic analysis

Mammary glands were spread on slides, fixed with Carnoy’s fixative, and stained in a carmine alum solution overnight. Subsequent steps included dehydration and clearing; whole mounts of mammary glands were captured with a microscope, and the ductal distance and terminal buds measured using ImageJ software.

### Blood biochemical examination

Blood samples from L2d mice were centrifuged to obtain serum and IGF-1 and IGFBP3 levels in serum measured (WT and Ho, n = 5) using commercial kits (R&D Systems, Minneapolis MN) following manufacturer’s instructions.

### Transcriptome analysis

Mammary tissues from L2d mice (WT and Ho, n = 3) were sent to Applied Protein Technology (China) for RNA sequencing. Total RNA was extracted and sequenced on the NovaSeq 6000 platform. Clean reads were carefully filtered for in-depth downstream analysis. Subsequently, FeatureCounts (http://subread.sourceforge.net/) was utilized to precisely quantify the reads aligned to each gene. The fragments per kilobase of exon model per million mapped reads (FPKM) value for each gene was calculated based on the gene length. Finally, to identify DEG between groups, DESeq2 was used. Genes with |log2Fold Change| > 0.585 and nominal *P*-value < 0.05 were defined as DEGs. A detailed compilation of the RNA sequencing dataset is included in Table S1.

### Lipidomic analysis

Milk samples collected from L2d sows of WT (n = 7) and Ho (n = 6) genotypes were sent to Cosmos Wisdom (Hangzhou, China) for lipid extraction and mass spectrometry detection. A pooled quality control (QC) sample was prepared by mixing equal aliquots of all individual samples to monitor analytical stability and data quality throughout detection. Lipid fractions were extracted via ultrasonic treatment, and the supernatant containing lipid extracts was collected for subsequent analysis.

Chromatographic separation was performed on a high-performance liquid chromatography system equipped with a Thermo Accucore™ C18 HPLC column. The declustering potential and collision energy for each lipid class were optimized using corresponding reference standards, and all instrumental operations controlled using analyst 1.6 software. Data quality was systematically assessed via QC sample stability, coefficient of variation, and PCA. The relative abundance of each lipid metabolite was determined through peak area normalization.

DLMs were screened using a combined multivariate and univariate statistical strategy. Orthogonal partial least squares discriminant analysis was implemented with the ropls R package to calculate Variable Importance in Projection (VIP) values, and univariate analysis (t-test or Wilcoxon test) applied for significance testing. Lipids meeting the thresholds of VIP > 1, nominal *P*-value < 0.05, and fold change > 2 were defined as DLMs. The full lipidomic dataset is provided in Table S4. KEGG pathway enrichment analysis of DLMs was performed using MetaboAnalyst 6.0 (https://www.metaboanalyst.ca/).

### Integrated transcriptome and lipidome analysis in mouse mammary glands

Mouse mammary gland transcriptome data were previously generated and analyzed by Fang et al [24]. Then, DEGs and DLMs were input into MetaboAnalyst 6.0 to identify key KEGG-enriched pathways. A complete list of enriched pathways derived from the integrated analysis is available in Table S2. Pearson’s correlation analysis between DEGs and DLMs was performed; only pairs with absolute correlation coefficient |r| > 0.95 were retained. IGF-1 and DEGs in the correlation network were submitted to STRING 12.0. Proteins lacking predicted physical or functional associations with IGF-1 were excluded with only interactions supported by a valid STRING combined score retained for network construction. Correlation and PPI networks were visualized using Cytoscape (v3.7.2).

### Codon usage bias analysis

RSCU was calculated as the ratio of observed to expected synonymous codon frequency under uniform usage. RSCU values > 1 and < 1 indicate positive and negative codon bias, respectively. Cross-species comparison of the *IGF-1* c.258 locus was performed using coding sequences from *Mus musculus*, *Sus scrofa*, and *Homo sapiens* retrieved from the NCBI RefSeq database. All calculations were performed using CodonW (v1.4.2) [75] (http://codonw.sourceforge.net/).

### LPL activity quantification

LPL activity was detected using Lipoprotein lipase Assay Kit (A067-1-1, Nanjing Jiancheng Bioengineering Institute, China). According to manufacturer’s instructions with minor modifications for the mammary gland. Briefly, 0.1 g mouse mammary gland tissue (WT and Ho, n = 5) was rinsed with ice cold saline. Tissue fragments were incubated in phosphate buffer (pH 7.4) containing 3.9 μg mL ¹ heparin at 37°C for 45 min to recover functionally active LPL. The supernatant was then collected for an enzymatic assay performed at 37°C, and the absorbance read at 550 nm. LPL activity was normalized to tissue weight and expressed as U per gram of tissue. All samples were assayed in technical duplicates.

### Detection of phosphatidic acid production

Phosphatidic acid production was detected using a Phosphatidic Acid Assay Kit (273335, Abcam, UK) according to manufacturer’s instructions. Briefly, 0.1 g of homogenized mammary gland tissue (WT and Ho, n = 5) was homogenized in 1-mL phosphatidic acid assay buffer. Lipid extraction was performed by adding 1.25 mL of a chloroform-methanol-12N HCl mixture (2:4:0.1, v/v/v) to 1 mL of tissue homogenate. The mixture was vortexed for 30 s, followed by addition of 1.25 mL of 1 M NaCl and centrifugation at 3000 ×g for 10 min at room temperature. The lower organic layer containing solubilized lipids was collected, and chloroform evaporated completely in a vacuum oven at 37°C. Extracted lipids were then solubilized in 5% Triton X-100 solution and used for phosphatidic acid measurement

### Quantitative real-time PCR

Total RNA was isolated using a total RNA extraction kit (Omega, USA) and reverse transcribed into cDNA with the gDNA Removal Kit (AT311, TransGen, China). During qRT-PCR, the prepared reaction mixture contained cDNA, supermix, nuclease-free water, and specific primers. The housekeeping gene *GAPDH* was used for normalization, and the relative quantification of gene expression calculated using the 2-ΔΔCt method. For m A quantification at the *IGF-1* c.258 locus, MeRIP-qPCR was adapted from Liu et al. with minor modification [76]. Briefly, total RNA was fragmented using VAHTS 2× Frag/Prime Buffer (N402-01, Vazyme, China) at 94°C for 5 min and purified with an Oligo Clean & Concentrator (D4060, Zymo Research, Orange County CA). Then, 10% fragmented RNA was reserved as input. The remaining RNA was incubated with an anti-m A antibody (1:200, 68055, Proteintech, China) in IPP buffer (10 mM Tris-HCl, pH 7.4, 150 mM NaCl, 0.1% NP-40) containing an RNase inhibitor at 4°C for 2–4 h. Antibody–RNA complexes were captured with Protein A/G Magnetic Beads (Thermo Fisher Scientific) at 4°C for 2 h, washed four times with IPP buffer, and eluted by Proteinase K digestion at 55°C for 30 min. Then, the eluted RNA was purified using an Oligo Clean & Concentrator and reverse-transcribed for qPCR. Relative m A enrichment was determined by using the ΔΔCt method, normalized to input controls, and expressed as fold change relative to the WT. All primer sequences used in this study are provided in Table S5.

### Western Blot

Proteins in mammary gland or MECs (WT and Ho, n = 3) were sequentially separated, transferred onto a membrane, and blocked before incubating overnight with primary antibodies: IGF-1 (1:1000, ab9572, Abcam), Lpin1 (1:500, YN1951, Immounway, Plano TX), FASN (1:1000, YM8374, Immounway,), Pparg (1:500, YT3836, Immounway), GAPDH (1:1000, AM4300, Thermo Fisher) and β-actin (1:1000, PA1-183, Thermo Fisher). Subsequently, proteins were incubated with a secondary antibody conjugated to horseradish peroxidase, goat anti-rabbit (1:4000, A0208, Beyotime, China), to develop protein bands. The intensity of the protein bands was quantified using ImageJ software.

### Statistical Analysis

Each experiment was performed ≥3 times. All results are expressed as mean ±SEM. All statistical analyses of lipidomic data were computed based on relative abundances. For data following a normal distribution, analyses were conducted using t-tests and one- or two-way analysis of variance with GraphPad Prism (v9.5). A *P*-value < 0.05 was considered to indicate statistical significance. \**P < 0.05*, ** *P* < 0.01, *** *P* < 0.001.

## Ethical statement

All animal experimental procedures were approved by the Animal Ethics Committee of Jilin University (SY202401017 and SY202407011). At the end of the experimental period, mice were euthanized by CO_2_ inhalation, while pigs received an intravenous pentobarbital overdose under general anesthesia.

## Data availability

The raw transcriptome data generated in this study have been deposited in the GSA [77] at the National Genomics Data Center (NGDC), China National Center for Bioinformation (CNCB), under accession number CRA044828, and are publicly accessible at https://ngdc.cncb.ac.cn/gsa/s/65O7qTUT. The raw lipidome data have been deposited in the OMIX [78] archive of NGDC/CNCB under accession number OMIX017087, and are publicly accessible at https://share.cncb.ac.cn/VOgeUb8qYc/OMIX017087/. All datasets analysed in this study are included in this article and its supplementary materials (Tables S1–S4).

## CRediT author statement

**Shuo Zheng:** Investigation, Writing—original draft, Writing—review and editing. **Jiayuan Fang:** Resources, Software, Investigation, Writing—review and editing. **Yi Li:** Investigation, Data curation. **Xingyu Xiao:** Software, Formal analysis. **Tong Su:** Visualization, Methodology. **Qinchuan Lv:** Investigation, Visualization. **Yunyun Cheng:** Formal analysis, Supervision. **Yuanyuan Xu:** Investigation, Writing—review and editing. **Linlin Hao:** Funding acquisition, Investigation.

## Competing interests

The authors have declared no competing interests.

## Acknowledgments

This work was supported by the National Natural Science Foundation of China (Grant No.32372963) awarded to Linlin Hao.

## Supplementary material Supplement Figures legends

**Supplement Tables**

**Table S1 Transcriptome sequencing data of mouse mammary gland**

**Table S2 Integrated analysis data of transcriptome and lipidome in mouse mammary glands**

**Table S3 Trait data of Bama Pigs**

**Table S4 Lipidomics data of Bama Pig milk**

**Table S5 Primer Sequences**

---

**Figure S1.**
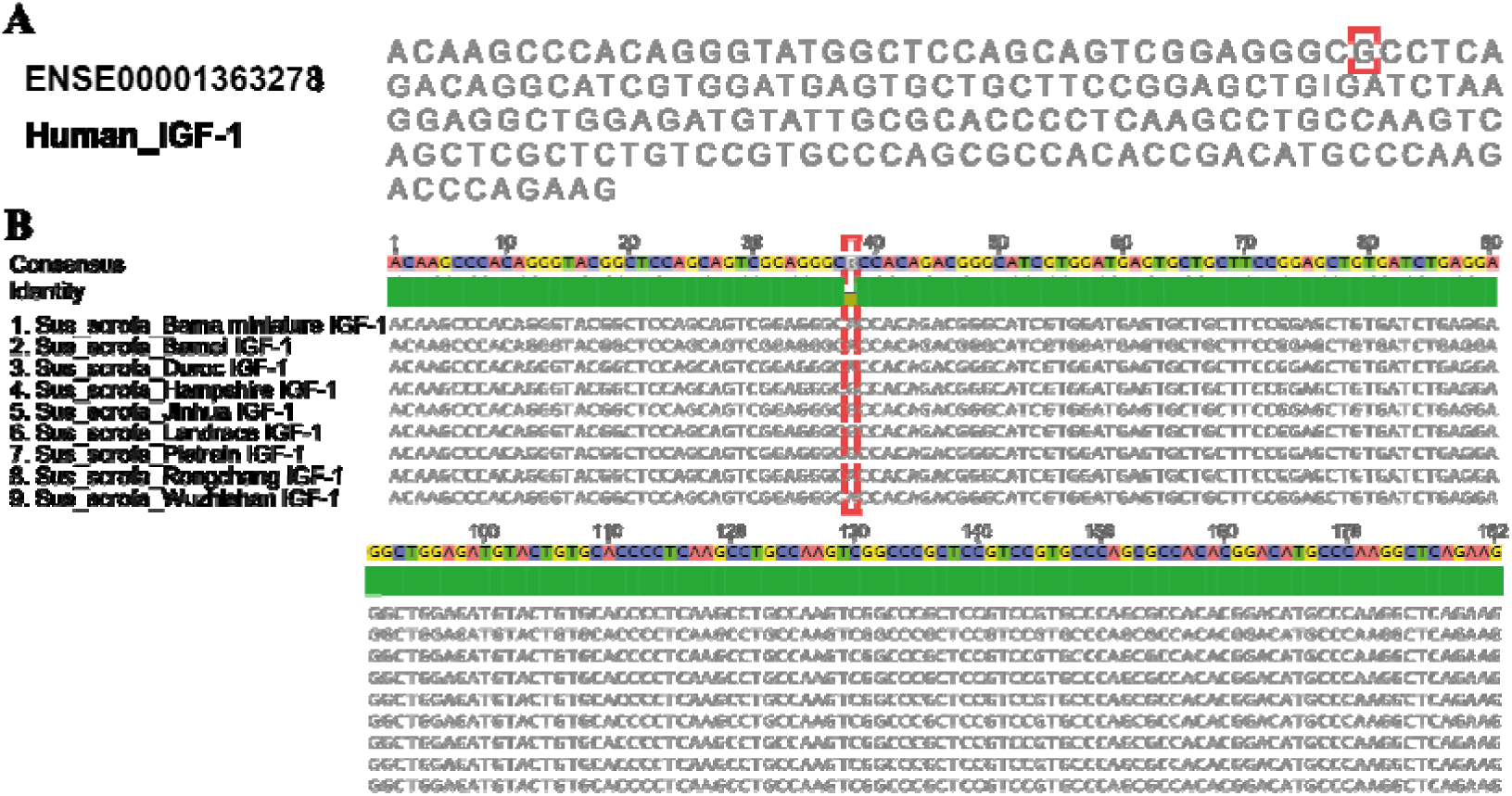
Genotypic variation of *IGF-1* c.258 in human and pig breeds. **A.** Partial coding sequence of the human *IGF-1*. **B.** Homology alignment across of *IGF-1* gene across pig breeds worldwide. All gene sequences were retrieved from the Ensembl database, and the c.258A>G variant has been confirmed to be polymorphic in human populations. The red box highlights the c.258A>G synonymous mutation locus.

**Figure S2.**
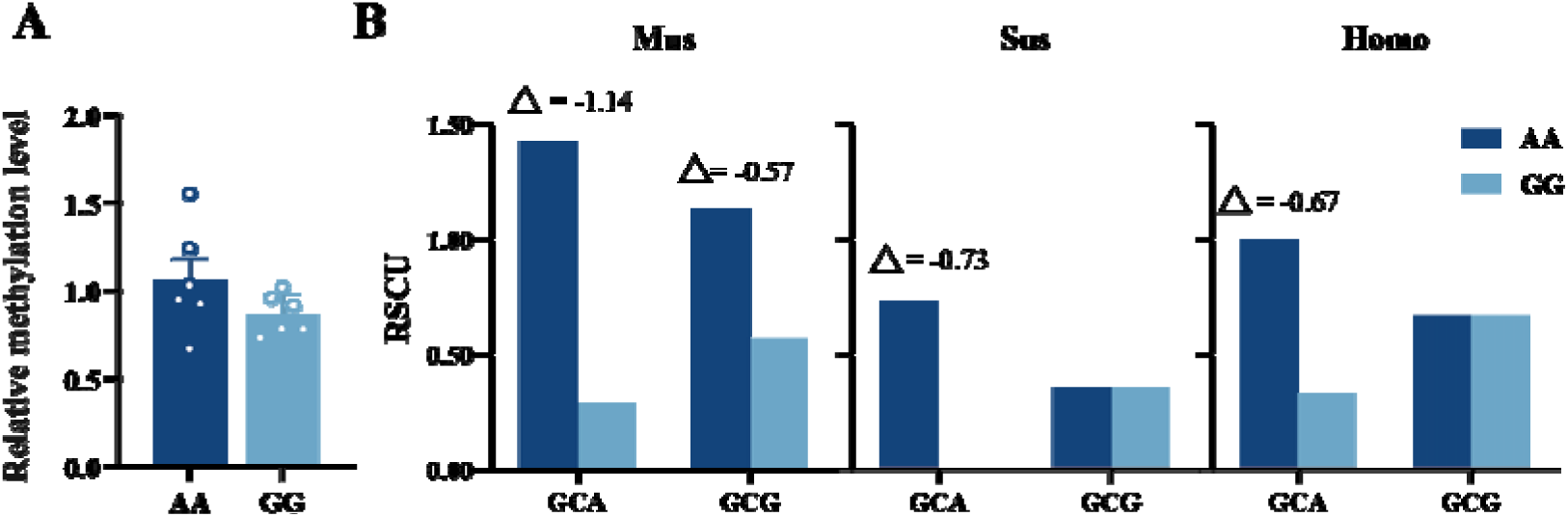
**m6A modification and cross-species codon usage bias at the *IGF-1*** **c.258 locus A.** Relative m6A methylation levels at the *IGF-1* c.258 locus in mammary glands of AA and GG genotype mice. **B.** RSCU, Relative synonymous codon usage analysis of GCA and GCG codons at the *IGF-1* c.258 locus in *Mus musculus*, *Sus scrofa* and *Homo sapiens*. △ denotes the difference in Relative Synonymous Codon Usage values between GCA and GCG. Data are presented as mean ± SEM; ns, not significant.

**Figure S3.**
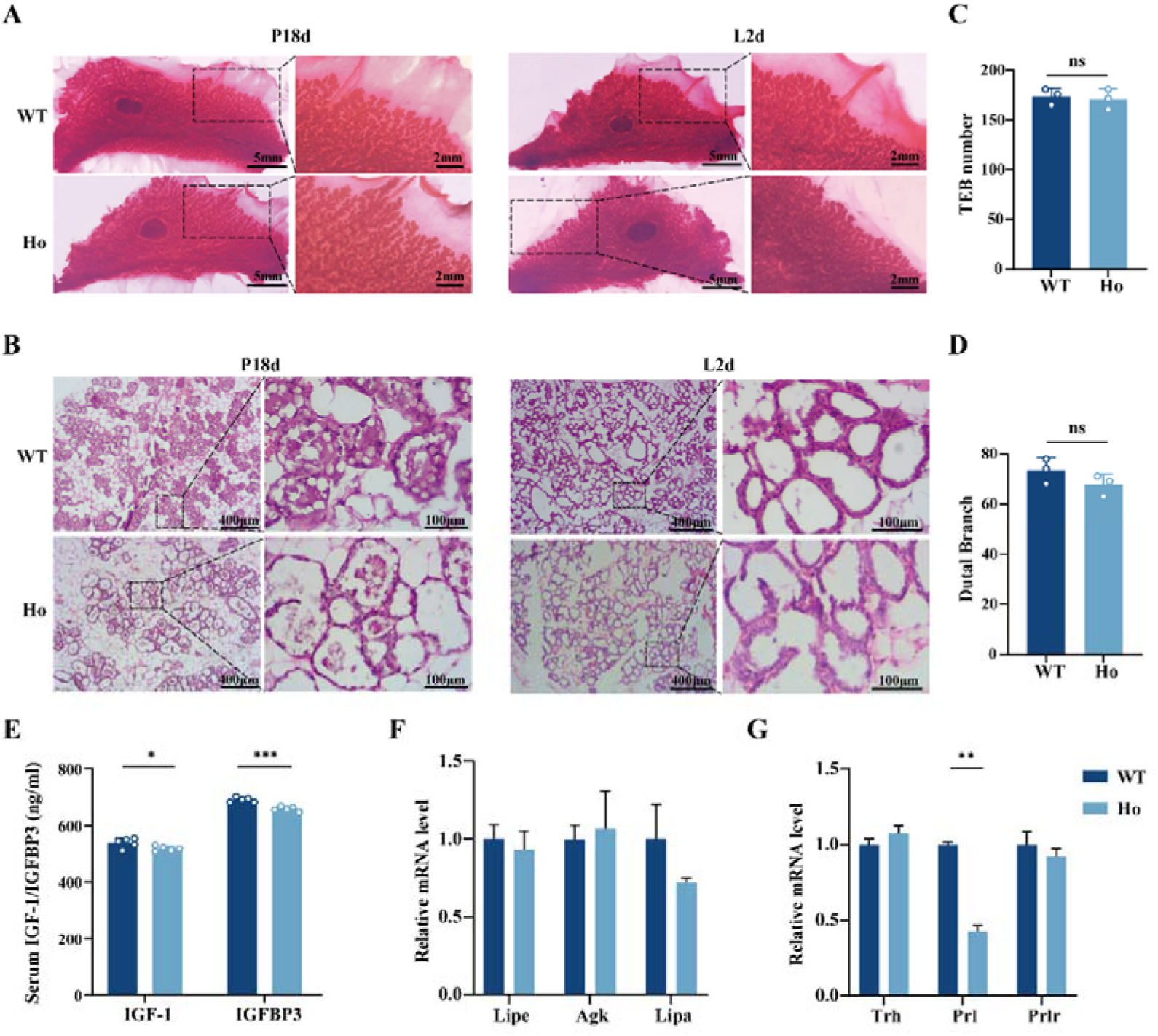
***IGF-1* c.258A>G does not affect the normal mammary gland development and systemic lipid supplement A.** Carmine-stained whole mount of mammary glands from WT and Ho mice at P18d, pregnancy day 18 and L2d. **B.** Hematoxylin and eosin staining of mammary glands from WT and Ho mice at P18d and L2d. **C. and D.** Statistics of mammary gland development, including the number of terminal end buds and dutal branch. **E.** Circulating IGF-1 and IGFBP3 levels in L2d mice. **F.** Expression of lipolysis related genes in liver of L2d mice. **G.** Expression of prolactin-related genes in pituitary gland of L2d mice. The data are presented as the mean ± SEM. \**P* < 0.05, ** *P* < 0.01, *** *P* < 0.001.

**Figure S4.**
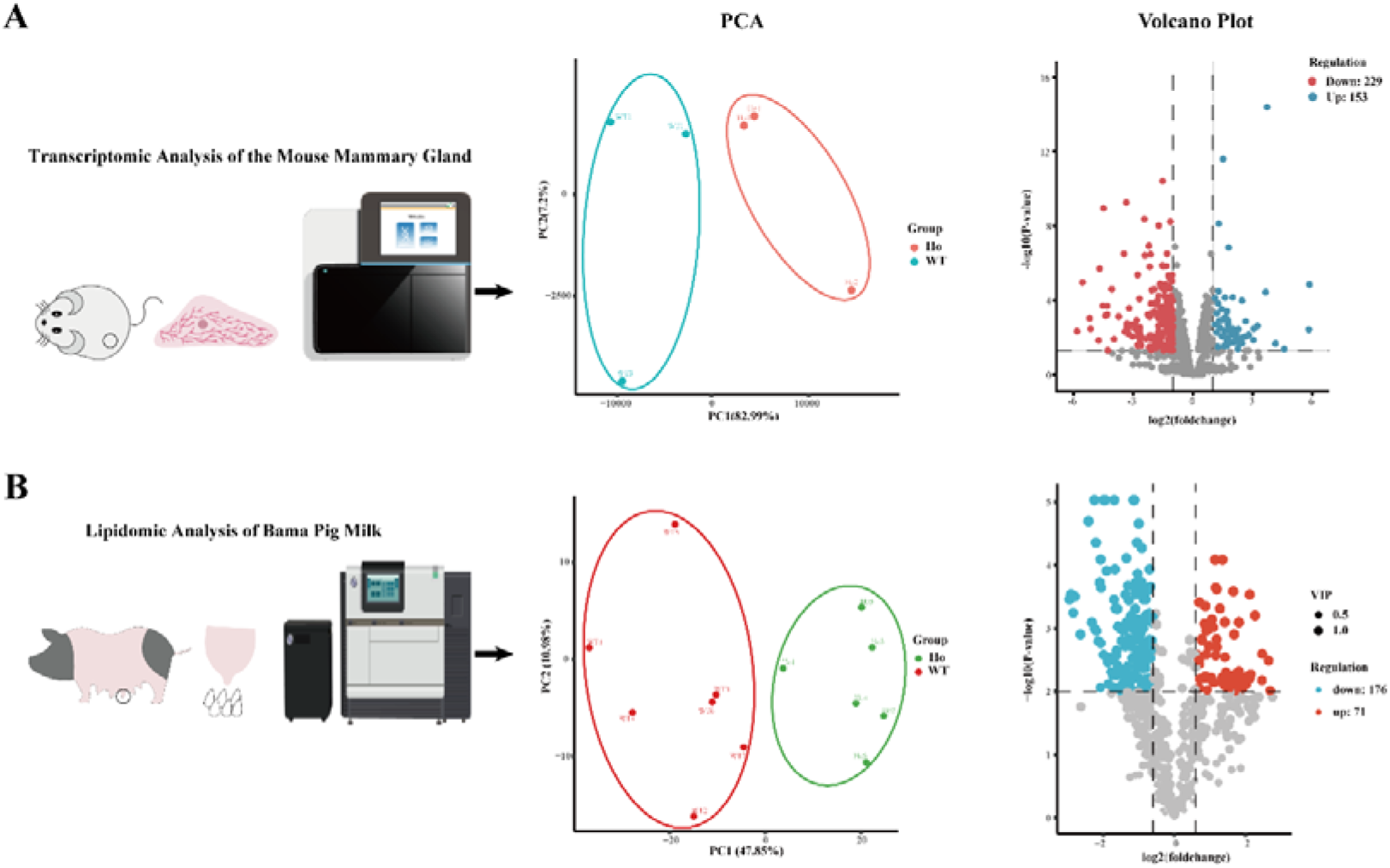
Flowcharts of multi-omics analyses. **A.** Transcriptomic analysis of mouse mammary gland. PCA, Principal component analysis clustering plots of inter-group genes (left) and volcano plots of DEGs (right). **B.** Lipidomic analysis of Bama pig milk. PCA clustering plots of inter-group lipids (left) and volcano plots of DLMs (right).

**Figure S5.**
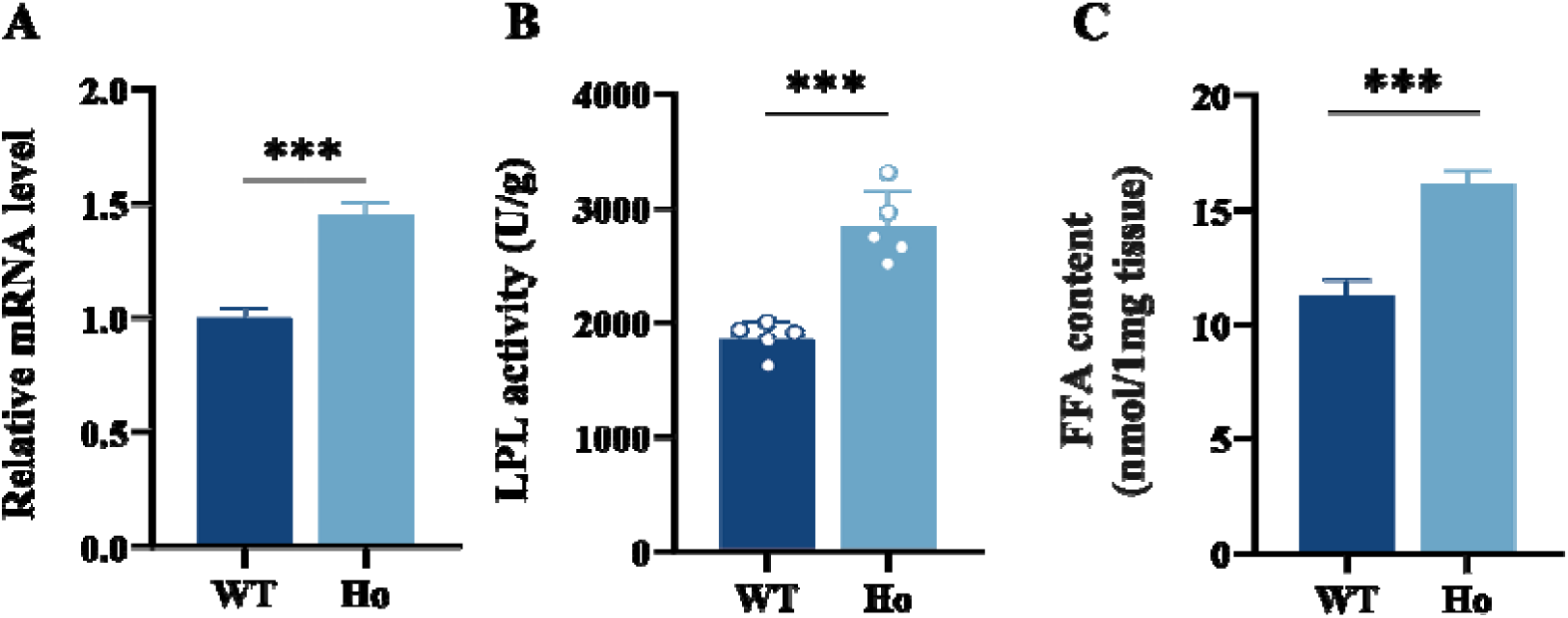
IGF-1 c.258A>G reduces LPL mRNA expression levels and enzymatic activity in the mammary gland A-C. LPL mRNA expression level (A), enzymatic activity (B) and free fatty acid content (C) in the mammary gland. The data are presented as the mean ± SEM. *** *P* < 0.001.

**Figure S6.**
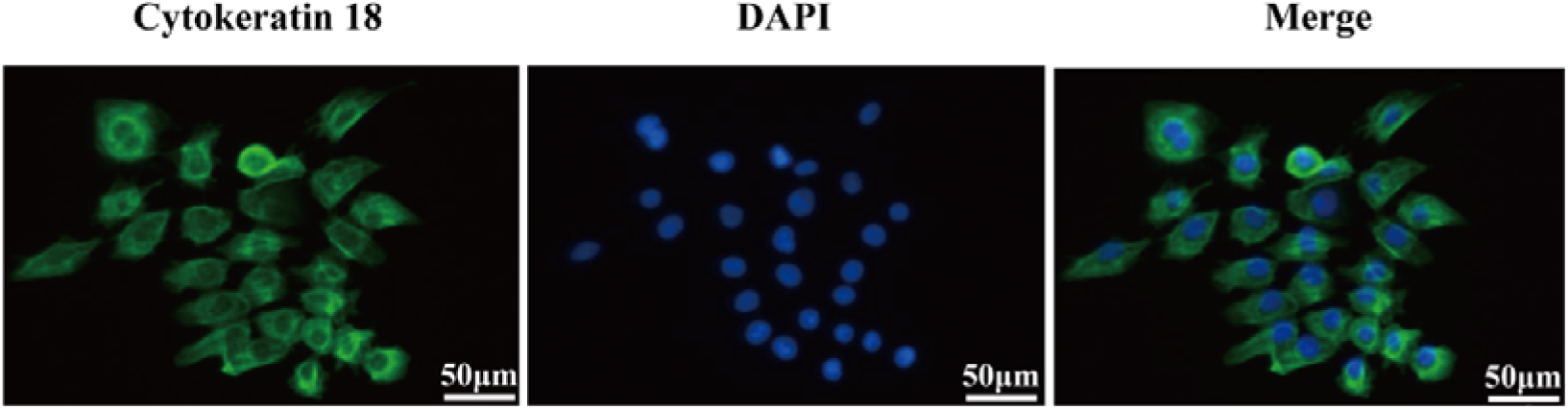
Identification of primary mouse mammary epithelial cells. Fluorescent staining of cytokeratin 18 for the identification of MECs.

**Figure S7.**
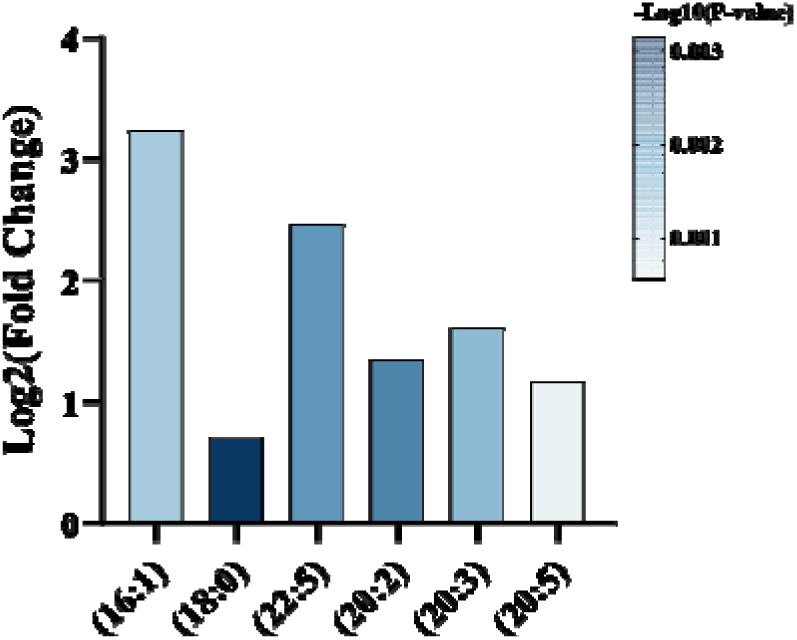
***IGF-1* c.258A>G alters FAs composition in sow milk** The intensity of individual fatty-acyl chains associated with different fatty acid classes. The transparency of each bar is proportional to the significance values, which are displayed as - log10 (*P*-value). The gray bars indicate those with *P* > 0.05.

**Figure S8.**
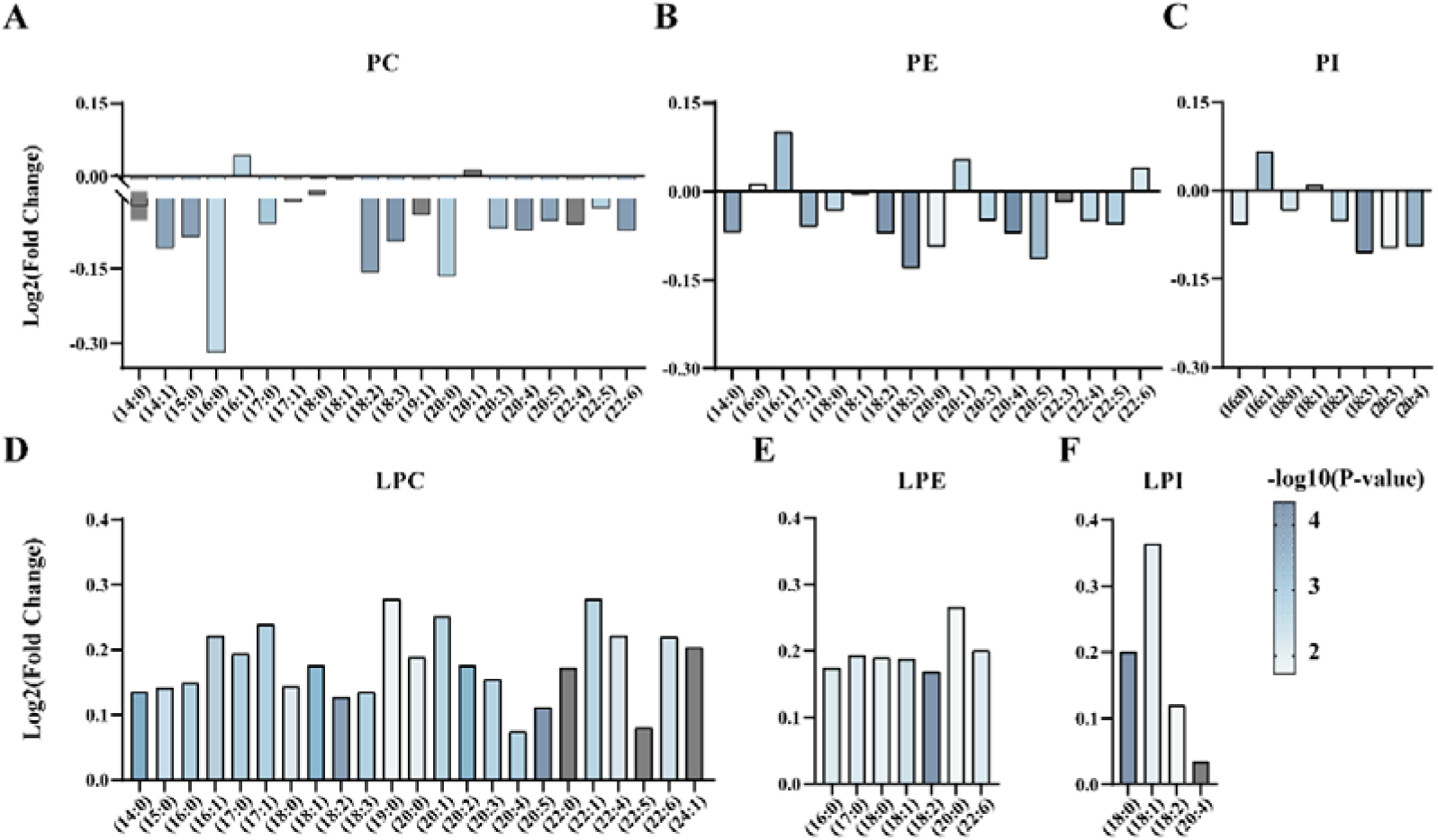
***IGF-1* c.258A>G alters glycerophospholipids composition in sow milk A-F.** The intensity of individual fatty-acyl chains associated with different glycerophospholipid classes. The transparency of each bar is proportional to the significance values, which are displayed as -log10 (*P*-value). The gray bars indicate those with *P* > 0.05.

**Figure S9.**
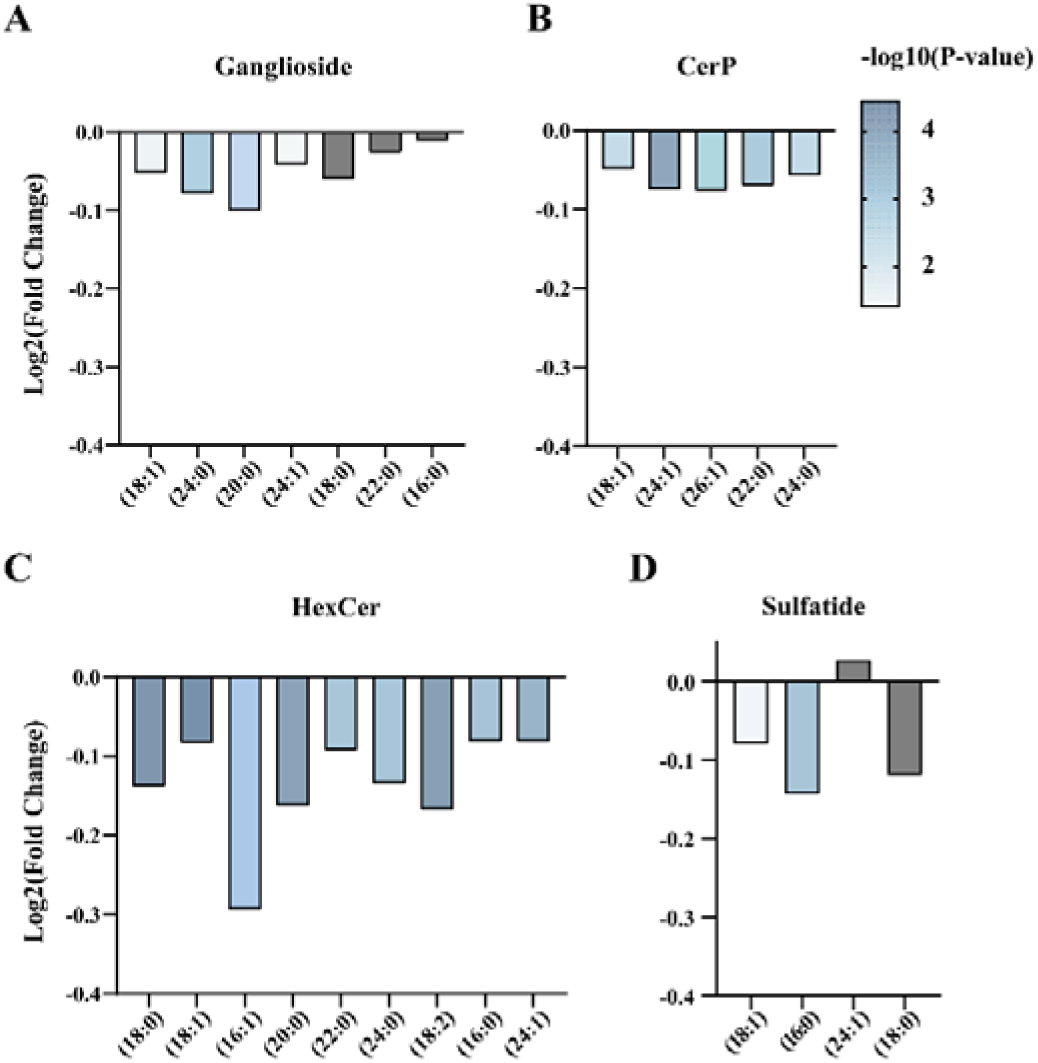
***IGF-1* c.258A>G alters sphingolipids composition A-D.** The intensity of individual fatty-acyl chains associated with different sphingolipids classes. The transparency of each bar is proportional to the significance values, which are displayed as -log10 (*P*-value). The gray bars indicate those with *P* > 0.05.

**Figure S10.**
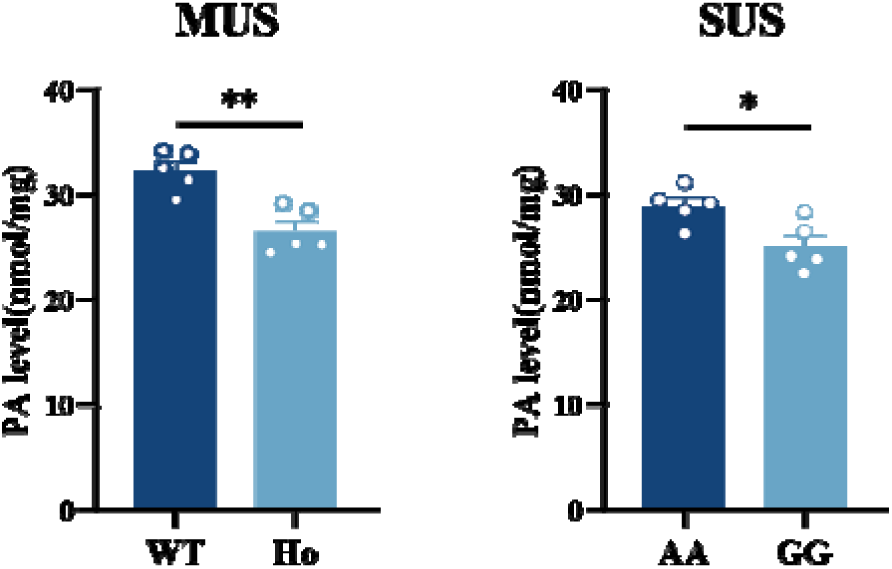
***IGF-1* c.258A>G reduces phosphatidic acid level** *IGF-1* c.258A>G reduces phosphatidic acid levels in the mammary glands of mice (left) and pigs (right). The data are presented as the mean ± SEM. \**P* < 0.05, ** *P* < 0.01.

**Figure S11.**
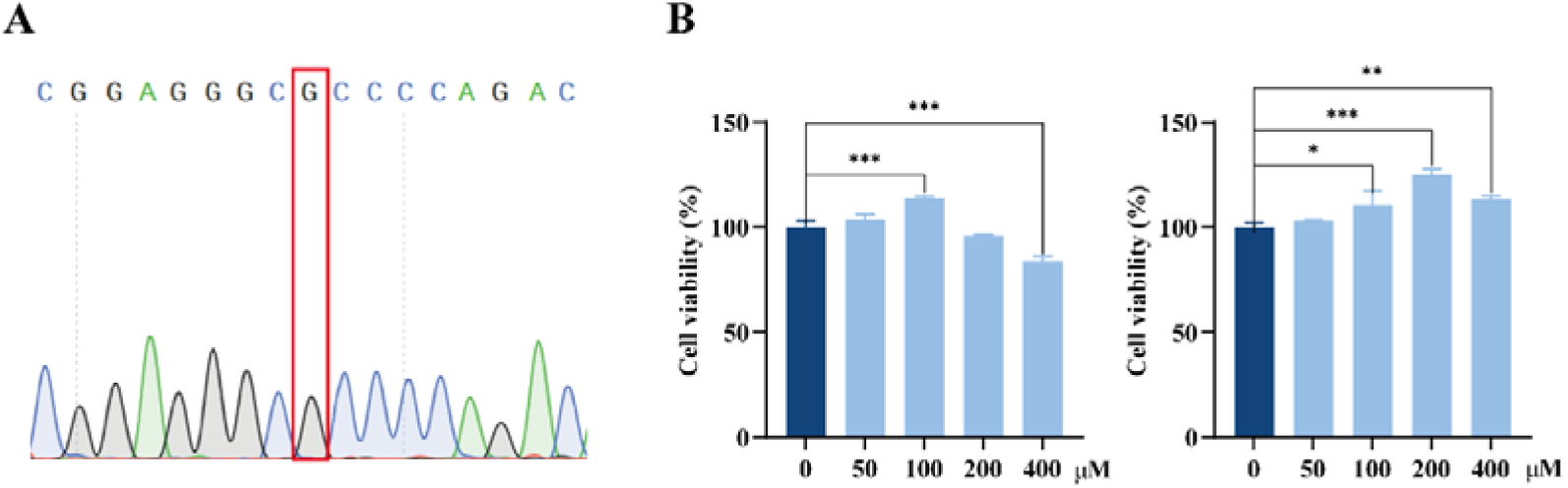
Genotyping of the *IGF-1* c.258 locus and determination of treatment concentrations of MCFAs in MCF-10A. **A.** Sequencing diagram of genotypes at *IGF-1* c.258 locus in MCF-10A. **B.** Proliferation assay of cells treated with lauric acid (left) and myristic acid (right) in MCF-10A. The data are presented as the mean ± SEM. \**P* < 0.05, \*\**P* < 0.01, \*\*\**P* < 0.001.

**Figure S12.**
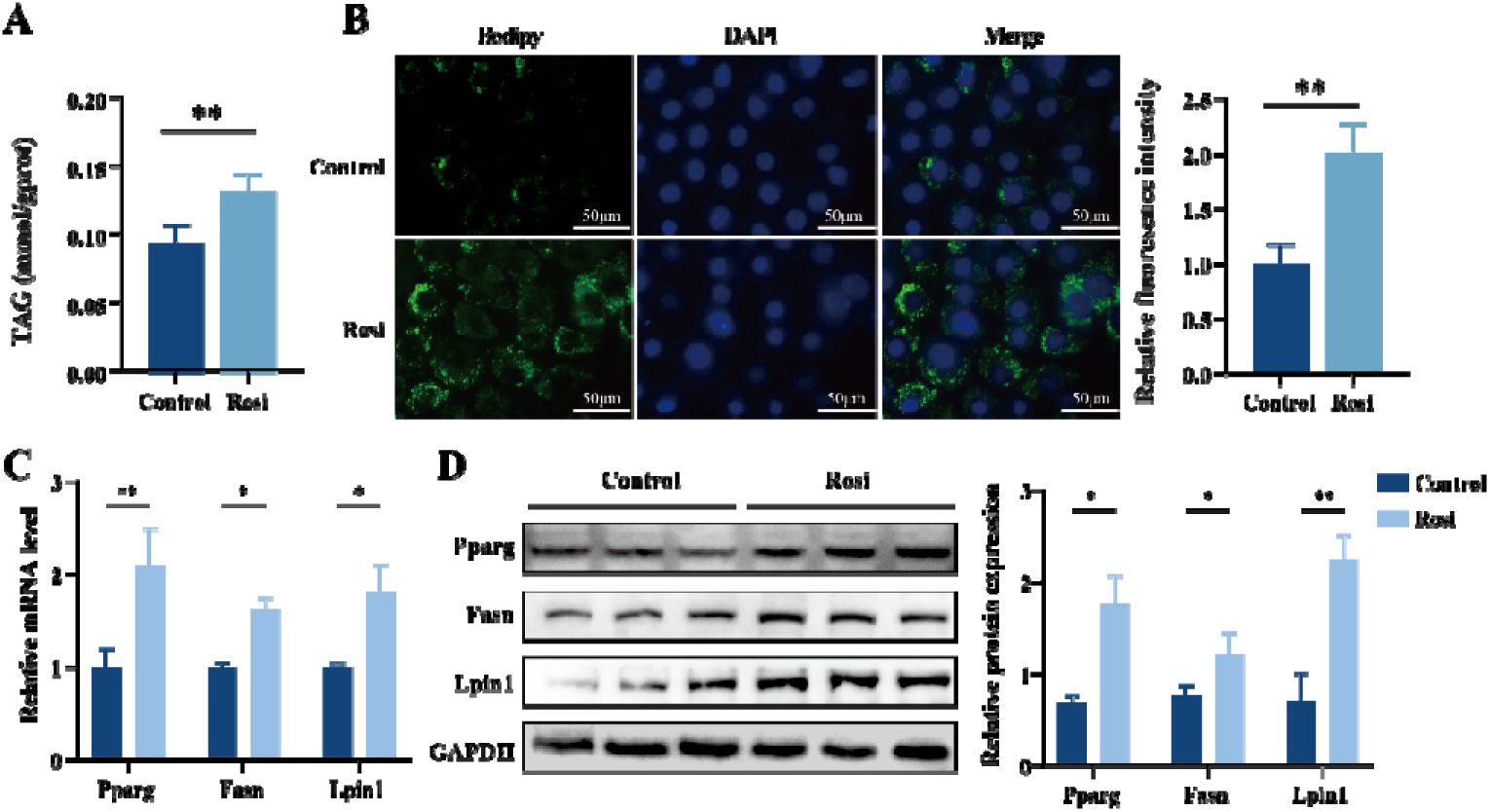
PPARG-FASN/LPIN axis regulates TAG synthesis in the mammary gland. **A.** TAG content in MCF-10A cells treated with Rosi. **B.** Bodipy staining of LDs (left) and relative fluorescence intensity (right) in MCF-10A cells treated with Rosi. **C.** The mRNA expression levels of genes in MCF-10A cells treated with Rosi. **D.** The expression of proteins MCF-10A cells treated with Rosi. The data are presented as the mean ± SEM. \**P* < 0.05, \*\**P* < 0.01, \*\*\**P* < 0.001.

**Figure S13.**
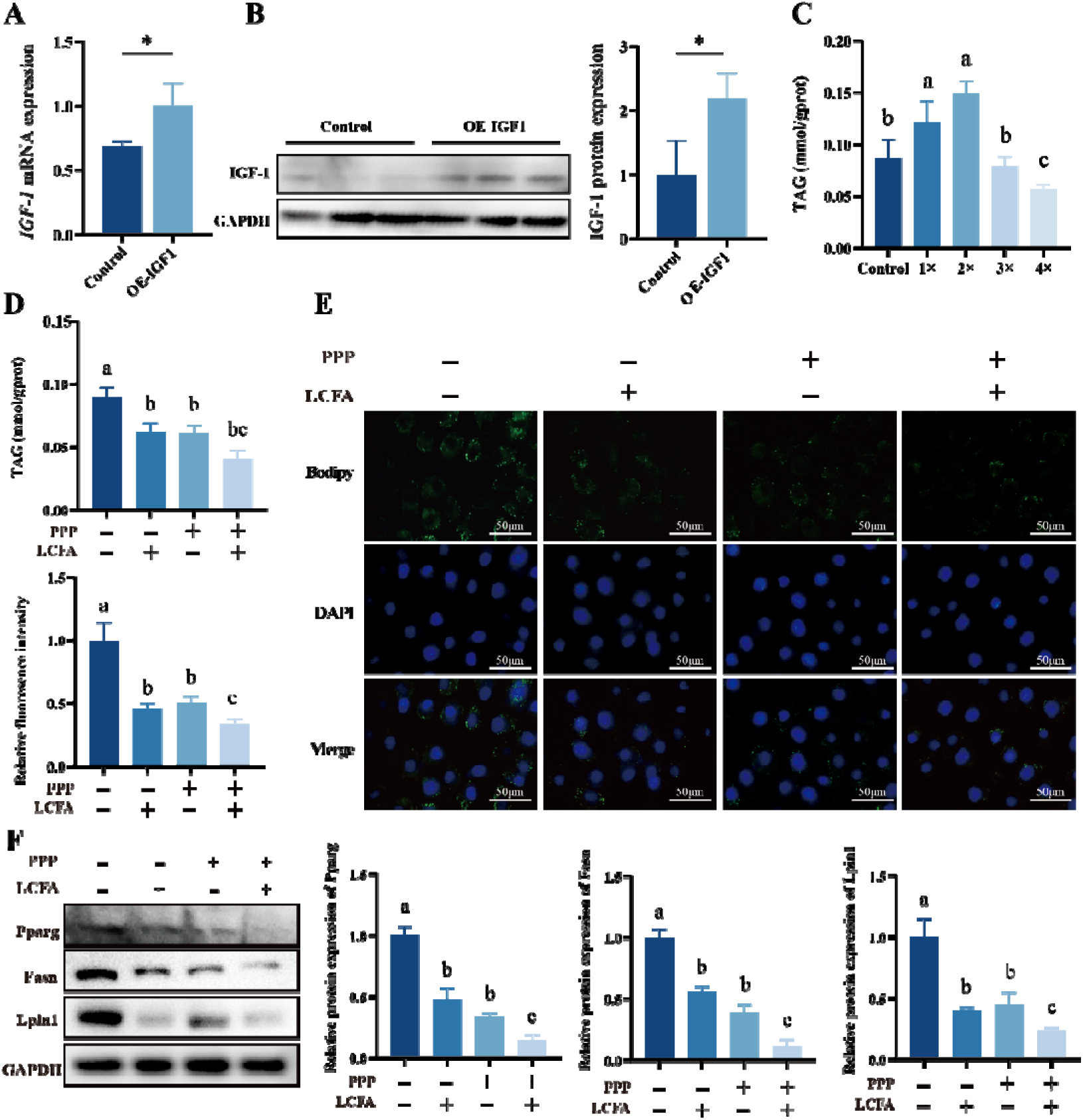
IGF-1 indirectly regulates PPARG-FASN/LPIN axis via the LPL-LCFA A. and. **B.** IGF-1 mRNA (A) and protein (B) expression analysis in MCFA-10A overexpressing IGF 1. **C.**TAG content in MCF-10A cells treated with gradient doses of an LCFA mixture. 1 × LCFA mixture = 100 μM oleic acid, 25 μM arachidonic acid and 8 μM α-linolenic acid. **D-F.** TAG content (D), bodipy staining of LDs (right) and relative fluorescence intensity (left) (E) and the protein expression levels of PPARG, FASN and LPIN1 (F) in MCF 10A cells treated with PPP and LCFA mixture. The data are presented as the mean ± SEM. \**P* < 0.05, Different letters indicate significant differences among groups (*P* < 0.05).

